# Structural basis for triacylglyceride extraction from mycobacterial inner membrane by MFS transporter Rv1410

**DOI:** 10.1101/2023.01.25.525346

**Authors:** Sille Remm, Dario De Vecchis, Jendrik Schöppe, Cedric A.J. Hutter, Imre Gonda, Michael Hohl, Simon Newstead, Lars V. Schäfer, Markus A. Seeger

## Abstract

*Mycobacterium tuberculosis* is protected from antibiotic therapy by a multi-layered hydrophobic cell envelope. Major facilitator superfamily (MFS) transporter Rv1410 and the periplasmic lipoprotein LprG are involved in transport of triacylglycerides (TAGs) that seal the mycomembrane. Here, we report a 2.7 Å structure of a mycobacterial Rv1410 homologue, which adopts an outward-facing conformation and exhibits unusual transmembrane helix 11 and 12 extensions that protrude ∼20 Å into the periplasm. A small, very hydrophobic cavity suitable for lipid transport is constricted by a unique and functionally important ion-lock likely involved in proton coupling. Combining mutational analyses and MD simulations, we propose that TAGs are extracted from the core of the inner membrane into the central cavity via lateral clefts present in the inward-facing conformation. The periplasmic helix extensions are crucial for lifting TAGs away from the membrane plane and channeling them into the lipid binding pocket of LprG.

## Introduction

*M. tuberculosis* (Mtb) is the notorious pathogen causing tuberculosis, a disease that claims ∼1.3 million lives annually world-wide^1^. The success of the pathogen depends on its hydrophobic multi-layered cell envelope which is both a formidable barrier offering protection from the host environment and an interface mediating host-pathogen interactions during infection. The mycobacterial cell wall is reminiscent of the cell envelope of Gram-negative bacteria in that Mtb has a periplasmic space that connects the inner membrane to an outer membrane^2^. However, the compositions of these compartments differ greatly^3,4^. In the ∼20-30 nm wide periplasmic space^2,5^, an arabinogalactan layer serves as a connector between the peptidoglycan layer and the outer membrane as it is covalently attached to both. The outer membrane, also called mycomembrane, is composed of two asymmetrical leaflets. The inner leaflet mainly consists of mycolic acids^6^, which are linked to the arabinogalactan via ester bonds while the outer leaflet is formed from various non-covalently bound lipid species^3,4^. At present, the exact composition and amount of the latter, is a subject of debate^3,4^.

Rv1410 (P55) is a major facilitator superfamily (MFS) transporter from Mtb. It is encoded in an operon together with lipoprotein LprG (Rv1411), which is embedded in the outer leaflet of the inner membrane via its lipid anchor^7^. The operon is highly conserved among mycobacterial species and both proteins are functionally contributing to intrinsic drug tolerance in mycobacteria^8^. Rv1410 has been initially described as a drug efflux pump^9–11^. However, more recent work has shown that Rv1410 indirectly contributes to drug tolerance by sealing the mycomembrane^8^ with triacylglycerides^12^.

Using lipidomics, the loss of the *lprG/rv1410c* operon or the *rv1410c* gene in Mtb was shown to result in intracellular accumulation of TAGs. Conversely, operon overexpression led to elevated secretion of TAGs into the culture medium, hence demonstrating that Rv1410 is a TAG transporter^12^. LprG has also been associated with the surface display of lipoarabinomannans, which can be tetraacylated or, akin to TAG, triacylated^13–15^. Crystal structures of LprG revealed a hydrophobic pocket that is able to accommodate lipids with two or three alkyl chains^12,13^ and *in vitr*o experiments with purified non-acylated LprG that lacks its lipid anchor show that it is capable of transferring TAGs between lipid vesicles^12^.

In mycobacteria, TAGs can be synthesized by several diacylglycerol acyltransferases^16,17^ of which many have been shown to localize to the inner membrane^18^. There, TAGs form lipid bodies and/or are transported to the outer membrane^19^. It is a plausible speculation that Rv1410 couples proton-motive force to active extraction of TAGs from the inner membrane to deliver them to the hydrophobic pocket of LprG. The molecular mechanism by which Rv1410 and LprG synergize to transport TAGs across the inner membrane and finally towards the mycomembrane is unclear. An LprG cross-linking study in live *Mycobacterium smegmatis* cells did not demonstrate a reproducible interaction between LprG and Rv1410^20^ and attempts to show complex formation *in vitro* with purified proteins were fruitless^8^. Nevertheless, it is evident from several studies that for optimal functionality, both proteins working in concert are needed^8,21,22^.

To address the questions of how Rv1410 transports TAGs and interacts with LprG, we determined the crystal structure of MHAS2168, an Rv1410 homologue from *Mycobacterium hassiacum*, in complex with an alpaca-raised nanobody. Based on the structure, we conducted molecular dynamics simulations and an extensive mutational analysis in both Rv1410 and MHAS2168. We found structural features essential for lipid transport, enabling us to propose a model describing the mechanism of TAG transport by Rv1410.

## Results

### Determination of MHAS2168 structure

To understand the lipid transport mechanism of Rv1410, we aimed to solve its structure. While we were unable to obtain crystals of Rv1410, we succeeded to crystallize its close homologue MHAS2168 (sequence identity of 62%) from the thermophilic mycobacterial species *M. hassiacum* in complex with an alpaca-derived nanobody Nb_H2 using crystallization in lipidic cubic phase (LCP). Native datasets diffracting up to 2.7 Å were obtained (Table S1), but initial attempts to solve the structure by molecular replacement, using other MFS transporters as search models, failed. Therefore, we used the megabody (Mb) approach^23^ to enlarge Nb_H2 and determined a cryo-EM structure of the MHAS2168-Mb_H2 complex, resulting in a density map with an average resolution of 4 Å (Fig. 1a; Extended Data Fig. 1b; Table S2). In parallel, we analysed Rv1410 in complex with another megabody (Mb_F7) by cryo-EM, yielding a map of 7.5Å (Extended Data Fig. 1a; Table S2). The MHAS2168-Mb_H2 map enabled us to build an initial model with assigned side chains for the bulk of the transporter. Intriguingly, we detected a non-proteinaceous density that could correspond to a triacylated lipid at the transporter’s side wall (Fig. 1a). The initial transporter model was then used to phase the native 2.7 Å data obtained by LCP crystallization by molecular replacement. The asymmetric unit comprises two transporter/nanobody complexes (Extended Data Fig. 1d). Complete models for the transporter and the nanobody were built with good refinement statistics and geometry (Extended Data Fig. 1c; Table S1). In the following analysis, chains A (MHAS2168) and B (Nb_H2) are used due to better map quality. In fact, the electron densities for the backside of Nb_H2 in chain D were blurry and explain the comparatively poor RSRZ scores of the model (Table S1).

**Figure 1.**
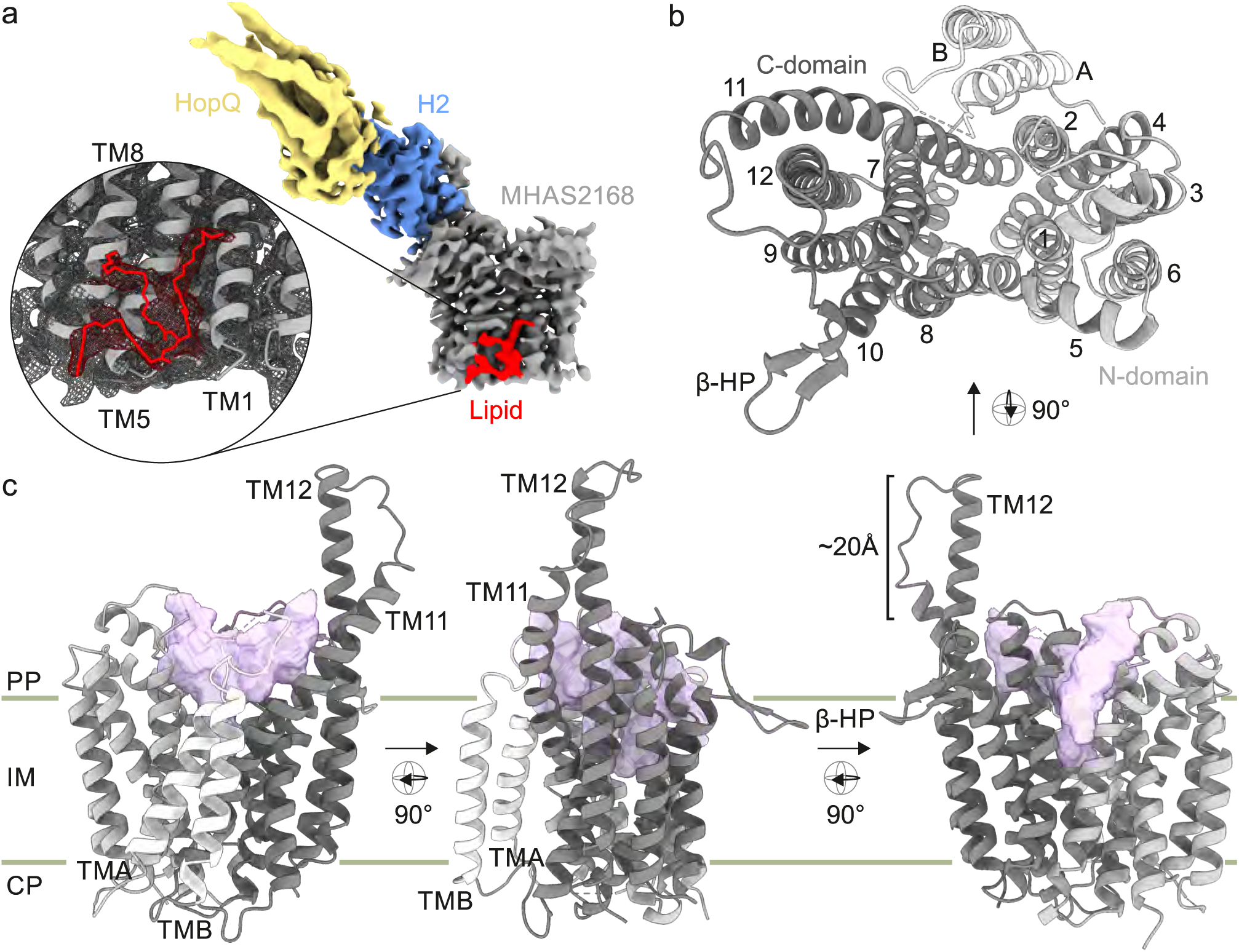
Architecture of MHAS2168. (a) 4 Å cryo-EM map of MHAS2168-Mb_H2 complex. MHAS2168 (gray), nanobody H2 (blue), megabody HopQ domain (yellow), non-proteinaceous density that likely corresponds to a triacylated lipid (red). Inset: The TAG species tripalmitoylglycerol fitted into the non-proteinaceous density. (b) Periplasmic top view of MHAS2168 2.7 Å crystal structure (nanobody not depicted). N-domain, transmembrane helices 1-6-light gray. Linker helices A and B-white. C-domain, transmembrane helices 7-12-dark gray. (c) Side views of MHAS2168 2.7 Å crystal structure. Color scheme same as in (b). Central cavity volume-light purple. β-HP - β-hairpin; IM -inner membrane; PP -periplasm; CP -cytoplasm; TM-transmembranehelix.

### MHAS2168 architecture

Both cryo-EM and crystal structures show MHAS2168 in its outward-facing (OF) conformation (Fig. 1). MHAS2168 adopts a canonical MFS transporter fold, featuring an N- and a C-domain each composed of six transmembrane helices (TMs). Since Rv1410 and MHAS2168 belong to the drug:H^+^ antiporter-2 (DHA2) subclass of MFS transporters (http://www.tcdb.org), they possess additional transmembrane linker helices A and B (TMA and TMB) between TM6 and TM7. The linker helices form a hairpin (Fig. 1b,c), such as seen in the structure of NorC^24^, as opposed to most other 14-helix MFS transporters whose linker helices are commonly arranged in a broader A-shape^25–28^. A striking element of MHAS2168 is a ∼20 Å long extension of TM11 and TM12 into the periplasm where we hypothesized the presence of a functionally important periplasmic loop in our previous work^8^. TM12 extends into the periplasm by 4 α-helical turns and is connected to the 2.5 α-helical turns-extended TM11 via a linker loop. Extended TM11 and TM12 seem to be a distinct feature of Rv1410 and its homologues, according to ColabFold^29^ structure predictions (Extended Data Fig. 2) and the crude 7.5 Å cryo-EM map of Rv1410-Mb_F7 complex (Extended Data Fig. 1a). As a last conspicuous structural element, MHAS2168 features a small extracellular β-hairpin between TM9 and TM10 (Fig. 1b,c), which extends along the membrane plane.

The ∼2300 Å^3^ outward-facing central cavity of MHAS2168 is well-equipped to accommodate lipids, in that the cavity walls of both domains are very hydrophobic, even compared to other MFS lipid transporters LtaA^30,31^ and MFSD2A^32–34^ (Extended Data Fig. 3). Curiously, the cavity does not extend as deeply into the transporter as in the case of other MFS lipid or drug transporters (Extended Data Fig. 3) due to a constricting salt bridge (comprised of E157 on TM5 and R426 on TM11) and its neighbouring residues in the cavity (Fig. 1c; Fig. 2a,b). There is a continuity between the central cavity and the membrane space, via narrow lateral openings between the N- and C-domains (Fig. 1b,c). The narrow lateral crevice between TM5 and TM8 is throughout its entire length 5-6Å wide, while the would-be lateral cleft between TM2 and TM11 widens from 4Å at the bottom to 12Å at the top. However, the TM2-TM11 cleft is shielded from the membrane by TMA and TMB. Therefore, it seems to be effectively blocked (Fig. 1b,c).

**Figure 2.**
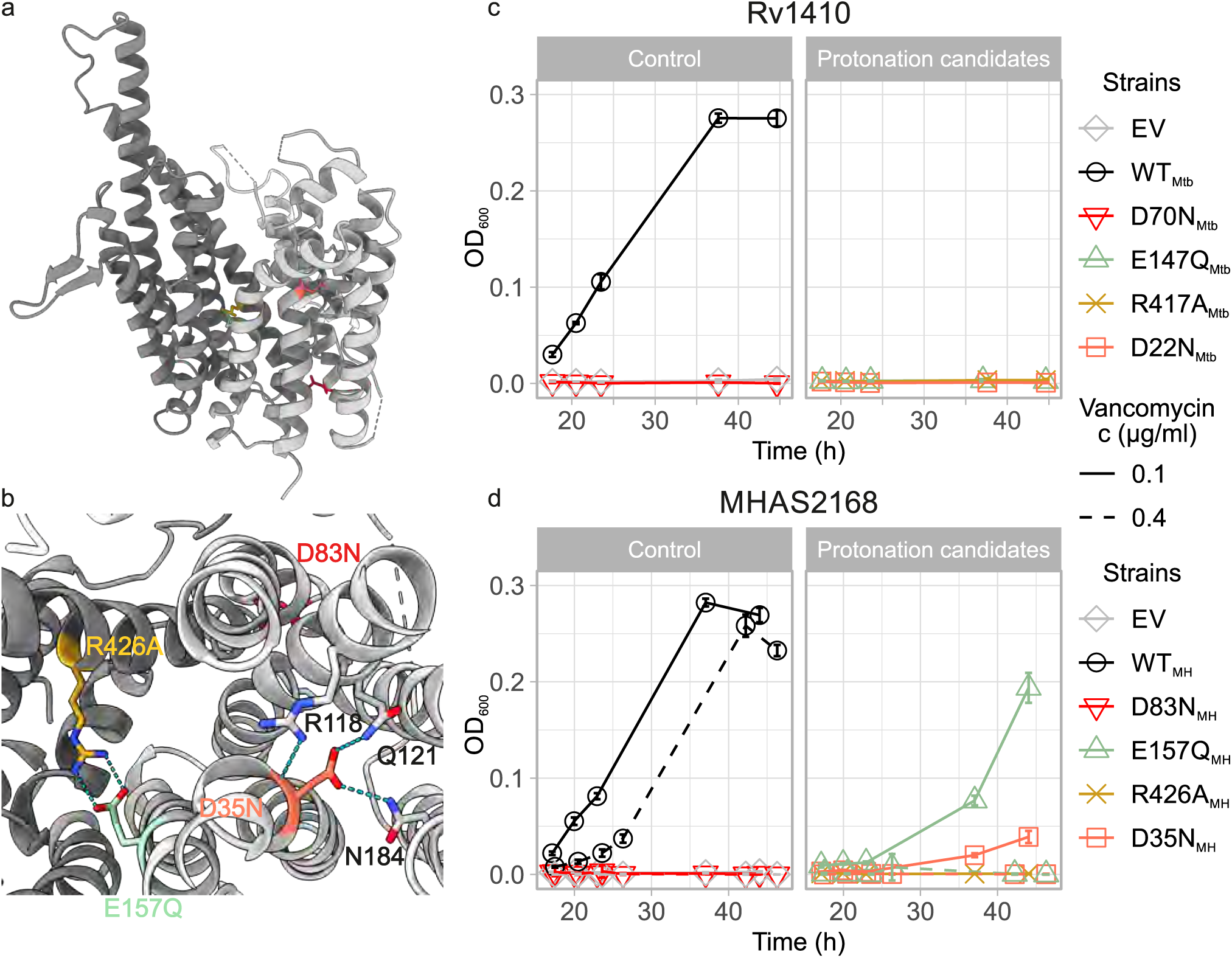
Two candidate loci for proton translocation. (a) Side view of MHAS2168. MHAS2168 color scheme same as **in** Fig lb. Point mutation sites are shown as colored sticks. (b) Enlarged periplasmic top view of MHAS2168 with mutated residues shown as colored sticks and interacting residues mentioned in the text as grey sticks, both with heteroatom depiction. Hydrogen bonds are shown as light blue dashed lines. (c) Vancomycin sensitivity assays in *M. smegmatis* dKO cells, complemented with empty vector control (EV), wild type LprG/Rv1410 operon (WT_Mtb),_ or mutant operons containing unaltered LprG (Rvl411) and mutated transporter Rv1410 as indicated. (d) Analogous analysis as in (c), with *M. smegmatis* dKO cells expressing instead wild type MHA S2167/68 operon (WTMH) or mutant operons containing unaltered LprG (MHAS2167) and mutated transporter MHAS2168 as indicated. The growth curves in (c) and (d) are representative of three biological replicates and error bars correspond to the standard deviation of four technical replicates.

### Experimental system to assess functionality of Rv1410 and MHAS2168 mutants in *M. smegmatis*

Rv1410 and LprG transport TAGs which secure the impermeability of the mycomembrane to certain drugs. Thus, the functionality of Rv1410 and LprG can be probed indirectly in *M. smegmatis* cells, by exploiting the change in cell envelope permeability, resulting in increased vancomycin influx to the periplasm if the *lprG/rv1410c* homologous operon *MSM3070/69* is deleted.^8^ This allows vancomycin to reach its periplasmic target, namely peptidoglycan precursors, and results in cell death. Complementation of the *M. smegmatis* deletion strain with *lprG/rv1410c* from Mtb or *MHAS2167/68* from *M. hassiacum* fully restores vancomycin resistance (Fig. 2c,d). Mutations in the transporter gene that cause loss of viability are therefore indicative of reduced transporter activity and TAG transport. Since complementation is carried out with the integrative pFLAG vector^35^ where the transporter is fused to a FLAG tag, protein production was confirmed by Western blotting for every mutant assessed in this work (Extended Data Fig. 4). As negative control, empty vector or motif A aspartate mutants of the MFS transporters which destabilize the outward-facing (OF) conformation^36,37^ (D70N_Mtb_ and D83N_MH_) were used for complementation (Fig. 2c,d).

### Two candidate loci for proton translocation

The only charged residues within the otherwise hydrophobic cavity of MHAS2168 are E157_MH_ and R426_MH_ which form a salt bridge (Fig. 2a,b) and constrict the bottom of the cavity. Curiously, this ion pair is conserved in 17 mycobacterial homologues of Rv1410 (Extended Data Fig. 5) but is absent from 46 previously characterized MFS transporters (Extended Data Fig. 6). The configuration of the glutamate as a possible proton acceptor/donor, the positively charged arginine to control pKa changes of the glutamate, and their location in the solvent-accessible cavity suggest a role of this ion pair in proton translocation^38^.

To test whether the ion lock residues contribute to energy coupling, E157_MH_ from MHAS2168 and the corresponding E147_Mtb_ from Rv1410 were mutated to glutamine. Similarly, R426_MH_ and R417_Mtb_ were mutated to alanine. When the Rv1410 salt bridge mutants were subjected to the vancomycin sensitivity assays at 0.1 µg/ml vancomycin, both E147Q_Mtb_ and R417A_Mtb_ mutations inactivated the transporter completely (Fig. 2c). Similarly, the R426A_MH_ mutant was unable to grow, but the E157Q_MH_ mutant showed completely abolished growth only under increased vancomycin stress (0.4 µg/ml) (Fig. 2d).

Another conserved carboxylate (Extended Data Fig. 5), D22 in Rv1410, has been shown to be required for transport before^21^. D22_Mtb_ and corresponding D35_MH_ are located in the middle of TM1 where their side chains point towards the center of the N-domain. D35_MH_ forms hydrogen bonds with Q121_MH_ (TM4, 1.8Å), N184_MH_ (TM6, 2.4Å), and R118_MH_ (TM4, 3.4Å) (Fig. 2b). D35_MH_ and the neighbouring R118_MH_ are potentially implicated in proton translocation due to analogously located aspartate-arginine salt bridges which were identified as the protonation/deprotonation sites via which substrate transport is coupled to proton translocation in certain sugar/H^+^ symporters^39–42^. In agreement with previously investigated mutants D22A_Mtb_ and D22E_Mtb_^21^, mutations D22N_Mtb_ and D35N_MH_ compromised the functionality of the transporter (Fig. 2c,d), suggesting that protonation/deprotonation of this carboxylate is necessary for transport function. While in the outward-open conformation of MHAS2168 the D35_MH_-R118_MH_ pair is not solvent-accessible, tunnels providing access to the bulk solvent might be present in other conformations. In summary, our data suggest that in Rv1410, both D22 and E147 are implicated in proton translocation and thus appear to play a key role in coupling the energy stored in the proton gradient to the efflux of TAGs.

### Molecular dynamics simulations suggest a mechanism of TAG extraction from the hydrophobic core of the membrane

To gain detailed molecular-level insights into how the transporter interacts with TAG substrate molecules, we performed coarse-grained molecular dynamics (MD) simulations of MHAS2168 in both outward-open (MHAS2168^OUT^; our crystal structure) and inward-open conformations (MHAS2168^IN^; homology model based on PepT_So2_) embedded in a phospholipid bilayer doped with TAGs, mimicking the mycobacterial plasma membrane (Fig. 3a; Table S3). The MD simulations revealed that TAGs are not fully embedded in the individual leaflets of the lipid bilayer, but rather segregate to the hydrophobic core of the membrane (Fig. 3b), likely owing to the absence of a charged head group. This observation has implications for potential transport mechanism of Rv1410, because TAGs need to be extracted from the membrane core, instead of being flipped from one leaflet to the other like phospholipids.

**Figure 3.**
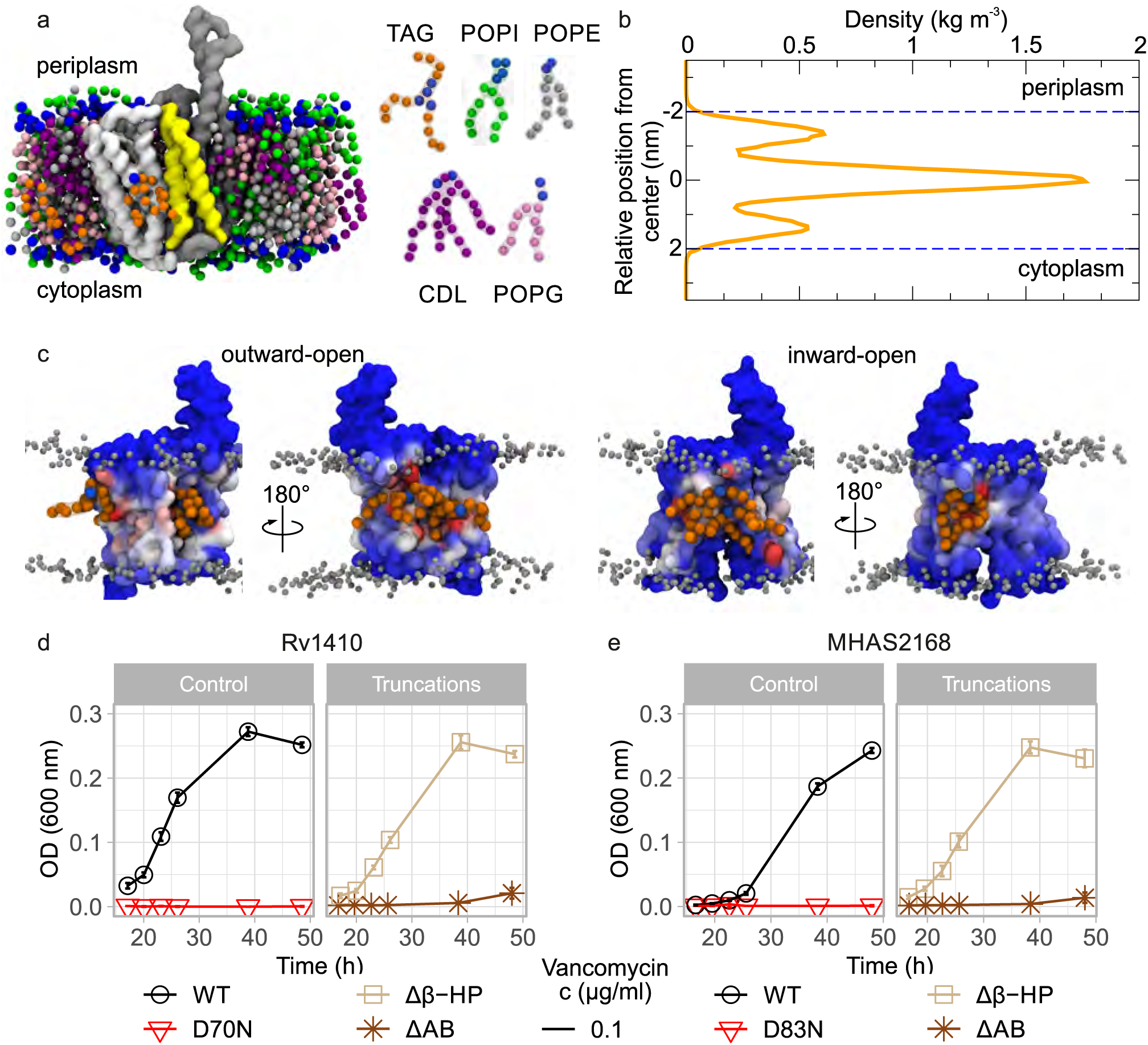
Molecular dynamics simulations-guided mutational analysis of MHAS2168. (a) Shown is the coarse-grained molecular dynamics simulation system of MHAS2168 in the outward-facingconformation. The mycobacterial plasma membrane lipid components are shown on the right, with headgroups colored blue. Solvent is not shown for clarity. (b) Density profile of TAGs along the membrane normal, centered with respect to the membrane midplane. The blue dotted lines are the average position of the phosphate headgroups. (c) Shown are the average number of TAG contacts from the MHAS2168^OUT^ (left) and MH AS2168^IN^ (right) simulations, respectively, projected on the surface of the protein and colored from blue (no contacts) to red (large number of contacts). (d) Vancomycin sensitivity assays in M. *smegmatis* dKO cells, complemented with wild type LprG/Rvl410 operon (WTMtb), or mutant operons containing unaltered LprG (Rv1411) and mutated transporter Rv1410 as indicated. (e) Analogous analysis as in (d), with *M. smegmatis* dKO cells expressing instead wild type MHAS2167/68 operon (WTMtt) ormutantoperonscontainingunalteredLprG(MHAS2167)andmutatedtransporterMHAS2168 as indicated. -ΔβHP - β-hairpin truncation; AB - truncation of linker helices Aand B. Thegrowth curves **in (d)** and (e) are representative of three biological replicates and error bars correspond to thestandard deviation of four technical replicates.

In the MD simulations, MHAS2168 strongly interacts with cardiolipin and phosphatidylinositol, which form an annular lipid belt around the transporter that shields it from other membrane components (Extended Data Fig. 7). The TAGs probe a range of different positions along the transporter TM helices, with preference for TM8 and TM10 (between the β-hairpin and TM5-TM8 lateral cleft) (Fig. 3c; Extended Data Fig. 7). Linker helices A and B and their surroundings form another hotspot for transporter-TAG interactions, especially in MHAS2168^IN^. To test the importance of these features to TAG transport, either TMA and TMB or the TM9-TM10 β-hairpin were truncated in Rv1410 and MHAS2168. Surprisingly, the loss of the β-hairpin did not affect lipid transport in neither Rv1410 nor MHAS2168, suggesting that this structural element plays a bystander role in the transport mechanism. In contrast, the deletion of the linker helices resulted in inactivation of both transporters, indicating that they are key to TAG transport (Fig. 3d,e).

### Mutational analysis indicates TAG entry into inward-facing cavity

Reflecting the hydrophobic nature of the TAG as substrate, the central cavity of outward-facing Rv1410 is particularly apolar (Extended Data Fig. 3). Its polarity was increased in a set of mutants by substituting a conserved leucine, which is situated in the middle of the hydrophobic C-domain cavity wall (Fig. 4c,d), with arginine or aspartate. Interestingly, only the L289R_Mtb_ and L299R_MH_ mutants were inactive; L289D_Mtb_ and L299D_MH_ mutants behaved like the wild type control strains at both lower (0.1 µg/ml) and higher (0.4 µg/ml) vancomycin concentrations (Fig. 4e,f). This discrepancy might be explained by the fact that arginines retain their charge in lipid and protein environments better than carboxylates^43^. The functional defect of arginine mutants suggests that TAGs enter the central cavities of Rv1410 and MHAS2168.

**Figure 4.**
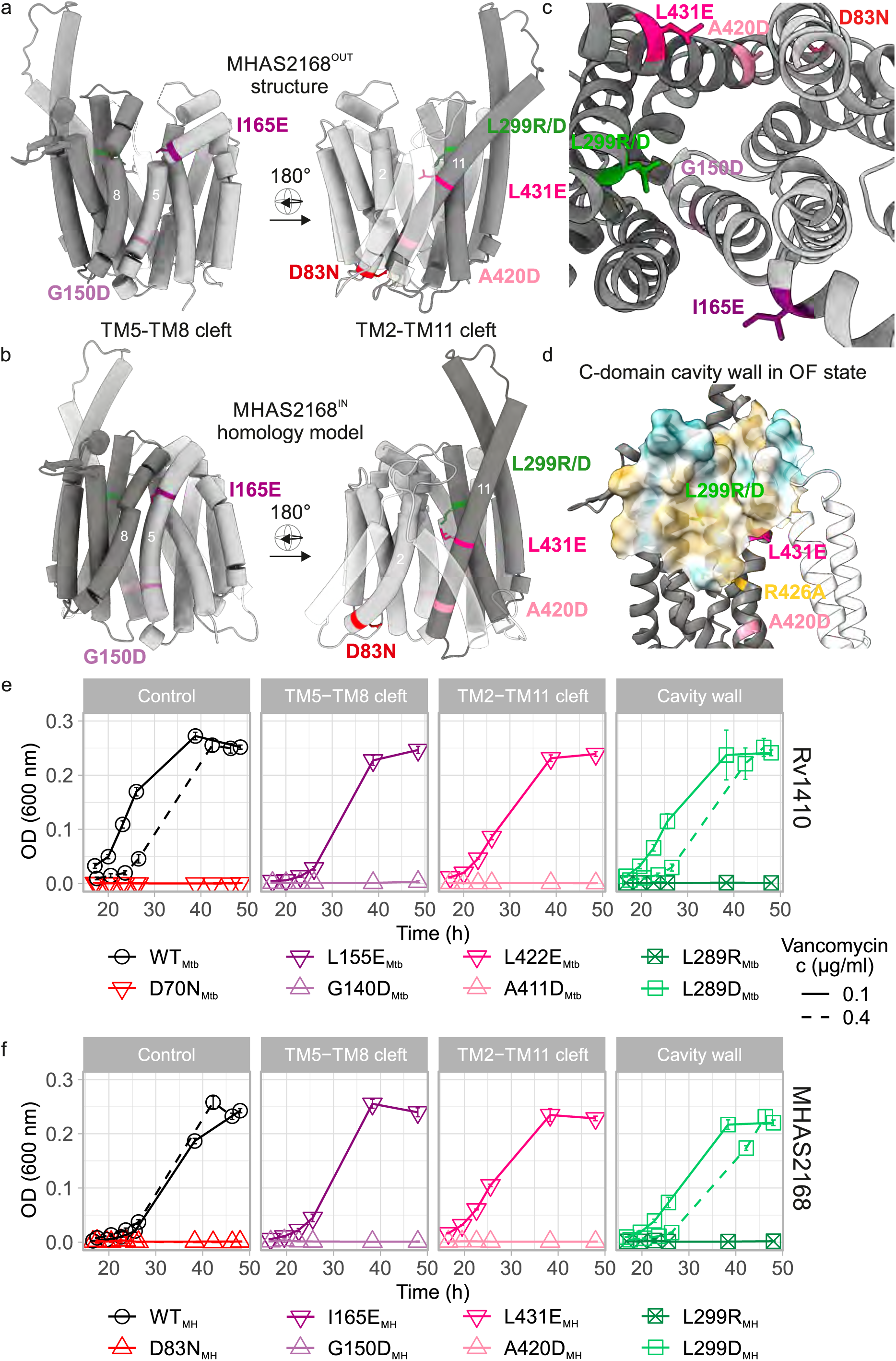
Potential TAG entry sites to the central cavity and cavity analysis. (a) Side view of lateral clefts between TM5-TM8 (left) and TM2-TM11 (right) in outward-facing conformation of MHAS2168 crystal structure. MHAS2168 color scheme same as in Fig 1b. Point mutation sites are shown as colored sticks. Linker helices A and B are depicted partially transparent. (b) Side view of lateral clefts between TM5-TM8 (left) and TM2-TM11 (right) in inward-facing conformation of MHAS2168 homology model. MHAS2168 and point mutation sites’ color scheme same as in panel (a). (c) Enlarged periplasmic top view of the central cavity and lateral clefts in outward-facing conformation of MHAS2168 crystal structure. MHAS2168 and point mutation sites’ color scheme same as in panel (a). (d) Hydrophobicity surface of the C-domain central cavity wall of MHAS2168 outward-facing crystal structure with mutated residues shown as colored sticks. Hydrophobicity color scheme: hydrophobic – gold; hydrophilic – cyan. (e) Vancomycin sensitivity assays in *M. smegmatis* dKO cells, complemented with wild type LprG/Rv1410 operon (WT_Mtb_), or mutant operons containing unaltered LprG (Rv1411) and mutated transporter Rv1410 as indicated. (f) Analogous analysis as in (e), with *M. smegmatis* dKO cells expressing instead wild type MHAS2167/68 operon (WT_MH_) or mutant operons containing unaltered LprG (MHAS2167) and mutated transporter MHAS2168 as indicated. The growth curves in (e) and (f) are representative of three biological replicates and error bars correspond to the standard deviation of four technical replicates.

According to our MD simulations, TAGs accumulate in the membrane core and therefore must enter the transporter through lateral openings between N- and C-domains lined with TM5 and TM8, or TM2 and TM11, respectively (Fig. 4a,b). To test whether the lateral crevices could serve as TAG entry or exit sites, residues in the middle of each lateral cleft were mutated into glutamates or aspartates to introduce polarity and block the opening (Fig. 4a,b,c; see Supplementary Information).

Functional assays (Fig. 4e,f) suggest that TAGs enter the cavity through both TM5-TM8 and TM2-TM11 lateral openings in the inward-facing state, as the corresponding mutants G140D_Mtb_, G150D_MH_, A411D_Mtb_, and A420D_MH_ were inactive. Another line of evidence that supports this notion is the observation of non-proteinaceous density in the cryo-EM map of the MHAS2168-Mb_H2 complex that could correspond to a TAG molecule or another tri-acylated lipid (Fig. 1a). This density is positioned in close proximity to the TM5-TM8 lateral cleft, opening in the inward-facing conformation.

As expected due to the obstruction by linker helices, the TM2-TM11 lateral crevice in the OF conformation is not involved in lipid transport as the L422E_Mtb_ and L431E_MH_ mutants display comparable growth to the wild-type operons at both lower (0.1 µg/ml) and higher (0.4 µg/ml) vancomycin concentrations. Even though the narrow TM5-TM8 lateral cleft in the OF state is connecting the central cavity and membrane, it does not seem to play a role in TAG transport. The corresponding mutants L155E_Mtb_ and I165E_MH_ grow similar to the wild type control at 0.1 µg/ml vancomycin concentration (Fig. 4e,f).

### Periplasmic TM11 and TM12 extensions guide TAG loading into LprG

It is clear from the MHAS2168 structure (Fig. 1c; Fig. 5a) and structure predictions of its homologues (Extended Data Fig. 2) that the “periplasmic loop” truncations we previously investigated in Rv1410^8^ manifest in fact as helix truncations (Fig. 5a), resulting in a less extended TM12. The Rv1410 truncation mutants were reproduced in MHAS2168, utilizing structure predictions performed with ColabFold platform^29^ for rational mutant design (see Supplementary Information). The two TM11 and TM12 truncation mutants in MHAS2168 were predicted to have the same length as Truncation 1_Mtb_ and Truncation 2_Mtb_ mutants in Rv1410 and connected by loops of similar length (Fig. 5a). Both MHAS2168 truncation mutants failed to transport lipids, while the Truncation 1_Mtb_ mutant retained some of its activity and the Truncation 2_Mtb_ mutant did not (Fig. 5b,c). Thereby, the previously observed loss in function of the TAG transporter was reproduced.

**Figure 5.**
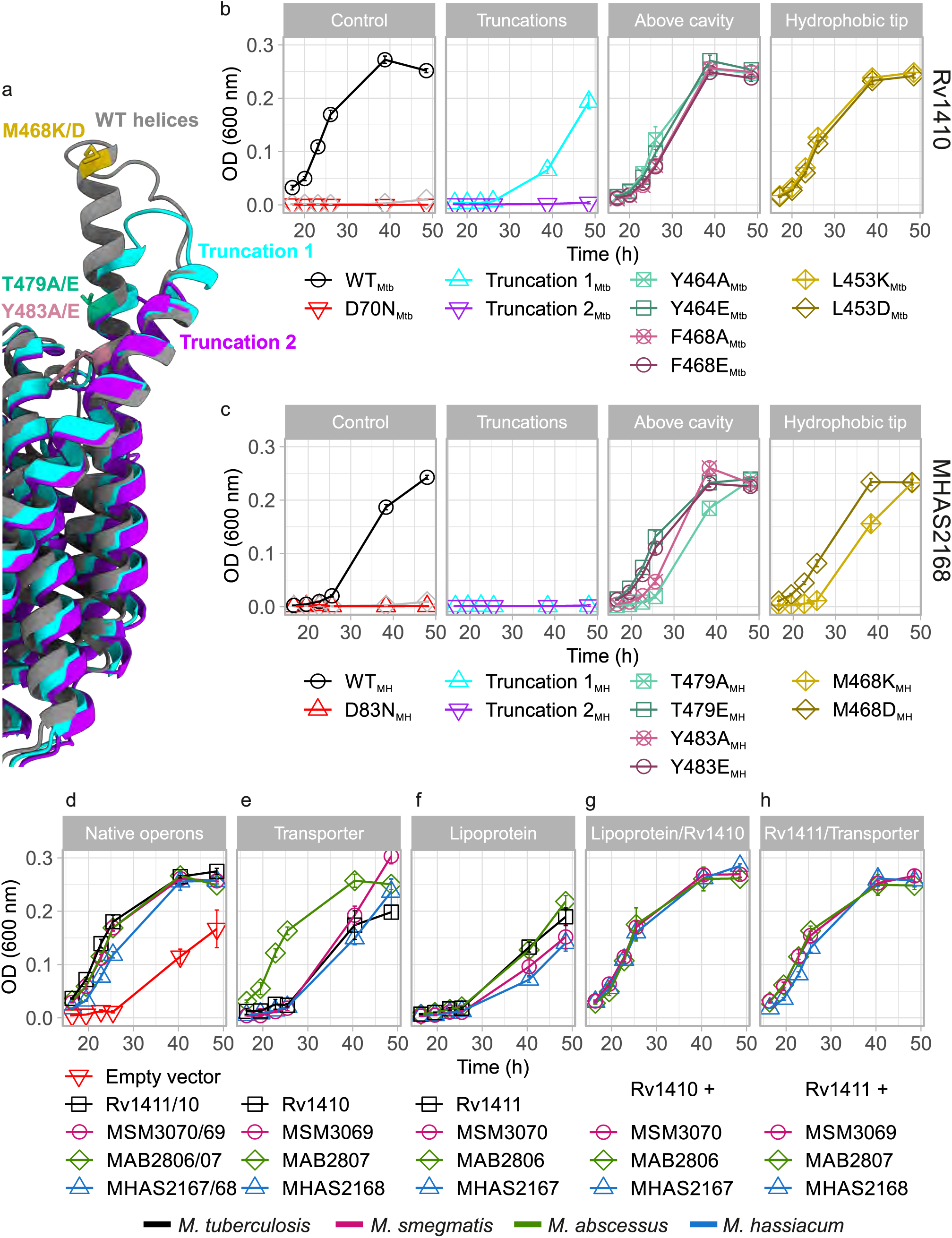
TM11 and TM12 periplasmic extensions and transporter-lipoprotein interactions. (a) Side view of MHAS2168^OUT^ crystal structure and predicted structures of truncation mutants 1 (purple) and 2 (light blue) of TM11-TM12 periplasmic extensions. MHAS2168 color scheme same as in Fig 1b. Point mutation sites are shown as colored sticks. (b) Vancomycin sensitivity assays in *M. smegmatis* dKO cells, complemented with wild type LprG/Rv1410 operon (WT_Mtb_), or mutant operons containing unaltered LprG (Rv1411) and mutated transporter Rv1410 as indicated. (c) Analogous analysis as in (b), with *M. smegmatis* dKO cells expressing instead wild type MHAS2167/68 operon (WT_MH_) or mutant operons containing unaltered LprG (MHAS2167) and mutated transporter MHAS2168 as indicated. (d) – (h) Vancomycin sensitivity assays in *M. smegmatis* dKO cells, complemented with different combinations of the transporter and/or lipoprotein from four mycobacterial species (*M. tuberculosis*; *M. smegmatis*; *M. abscessus*; *M. hassiacum*). (d) Complementation with native operons or empty vector. (e) Complementation with only the transporter. (f) Complementation with only the lipoprotein. (g) Complementation with a shuffled operon in which the transporter from *M. tuberculosis* (Rv1410) is accompanied by lipoprotein from the other three mycobacterial species. (h) Complementation with a shuffled operon in which the lipoprotein from *M. tuberculosis* (Rv1411) is accompanied by transporter from the other three mycobacterial species. Vancomycin sensitivity assays were carried out at 0.1 µg/ml concentration on panels (b) and (c) or at 0.08 µg/ml concentration on panels (d) – (h). The growth curves in (b) - (h) are representative of three biological replicates and error bars denote the standard deviation of four technical replicates.

An in-depth analysis of the MHAS2168 structure and the structure prediction models of its mycobacterial homologues revealed that although the primary structure of the TM11-TM12 extensions varies in both length (31-38 residues) and sequence, they share common features (Extended Data Fig. 2; Extended Data Fig. 5). Firstly, the TM12 length is very conserved (extra 4 α-helical turns) while there are small differences in the TM11 and TM11-TM12 loop lengths (Extended Data Fig. 2a). Secondly, hydrophobic patches are present on the side of the extensions facing the cavity (Extended Data Fig. 2b-d), and the residues on the tip of the TM12 extension (1^st^ α-turn) are commonly hydrophobic. Thirdly, aromatic residues are prevalent on TM12 above the cavity.

To study the functional role of these molecular features on the TM12 extension, single point mutations were generated with the aim to alter the biophysical properties of this region (Fig. 5a). Therefore, the aromatic or electroneutral residues of TM12 above the cavity were mutated either to alanines to remove phenol groups that might be involved in stacking interactions (Y464A_Mtb_, F468A_Mtb_, T479A_MH_, Y483A_MH_) or to glutamates to introduce charge and bulkiness to the side chains and thus likely disrupt lipid movement (Y464E_Mtb_, F468E_Mtb_, T479E_MH_, Y483E_MH_). Similarly, the hydrophobic residues at the tip of TM12 extension were mutated into lysines (L453K_Mtb_, M468K_MH_) or aspartates (L453D_Mtb_, M468D_MH_). While all of these mutants retained wild type activity in the vancomycin sensitivity assays under milder condition (0.1 µg/ml vancomycin; Fig. 5b,c), growth defects manifested in many of these mutant strains at higher vancomycin concentration (0.4 µg/ml; Extended Data Fig. 8c,d). In conclusion, the helical extensions cannot be removed without affecting the transporter’s function. Further, disruption of conserved molecular features on TM12 results in partially defective transport activity.

We hypothesized that the TM11 and TM12 periplasmic extensions might play a role in TAG exit from the central cavity by serving as an anchor point to place LprG into a favourable position for TAG loading. Although previous experiments failed to demonstrate physical interactions between purified Rv1410 and LprG^8^, transient low affinity interactions nevertheless may occur in the cellular context. We reasoned that if there was a specific physical interaction between the extended helices of Rv1410 and LprG, the operon partners might have co-adapted during evolution. Then, MFS transporters and lipoproteins from different mycobacterial species would not be able to act in concert. To examine this hypothesis, Rv1410 and LprG from *M. tuberculosis* were shuffled with their counterpart homologues from three mycobacterial species (namely *M. smegmatis*, *M. hassiacum*, and *M. abscessus*) without affecting the overall operon structure. *M. smegmatis* dKO complemented with any tested combination of transporter and lipoprotein was able to grow as fast as with the natively paired operons while complementation with the transporters (with the exception of MAB2807 from *M. abscessus*) or lipoproteins alone resulted in substantial growth defects (Fig. 5d-h). Considering the fact that protein-protein interactions are highly sensitive to small perturbations at the binding interface and the overall low degree of sequence conservation in the helical extensions, our data do not support specific transporter-lipoprotein interactions. Rather, our experiments suggest that transporter and lipoprotein functionally synergize to accomplish efficient TAG transport without the necessity to specifically interact at the protein level.

Alternatively, the importance of periplasmic helix extensions in TAG transport might lie in their interactions with the TAG during its transit from Rv1410 to LprG. Hence, we decided to study the dynamic process of TAG loading from the transporter into LprG directly. The ColabFold platform^29^ was used in multimer mode to build a complex of MHAS2168^OUT^ and LprG. All five obtained models consistently positioned LprG on the periplasmic side, with its hydrophobic cavity oriented towards TM11 and TM12 (Fig. 6; Extended Data Fig. 9a). A TAG molecule was inserted into the main cavity of the transporter and extended unbiased coarse-grained MD simulations of the MHAS2168^OUT^-LprG complex were carried out. Several events of TAG-loading into LprG were observed, at a rate of about one event per 100 μs of simulation time (Fig. 6). In addition to the loading of TAG into LprG, also the reverse unloading process was observed (Fig. 6a). TAG slides back from LprG into the transporter along the extended hydrophobic tunnel that is established upon the formation of the MHAS2168-LprG complex (Extended Data Video 1), suggesting a rather shallow energy landscape. Interestingly, in our simulations, the TAG molecule migrated to LprG only when two of its acyl tails were pointing up towards LprG, instead of only one (see Supplementary Information). The key residues of the transporter that line the hydrophobic tunnel and mediate the transitioning of TAG between the transporter and LprG are found in the loop connecting the linker helices A and B, TM 7 C-terminus, and particularly TM11 and TM12 (Fig. 6b,c; Extended Data Fig. 9; Table S4). For LprG, the residues that guide TAG loading cover its entrance mouth and form a broad hydrophobic surface that extends further into the hydrophobic cavity of LprG (Extended Data Fig. 9; Table S4). The simulation results are in line with a site-directed mutagenesis study showing that V91 in *M. tuberculosis* LprG, corresponding to T87 in *M. hassiacum* (Table S4), plays a key role^12^.

**Figure 6.**
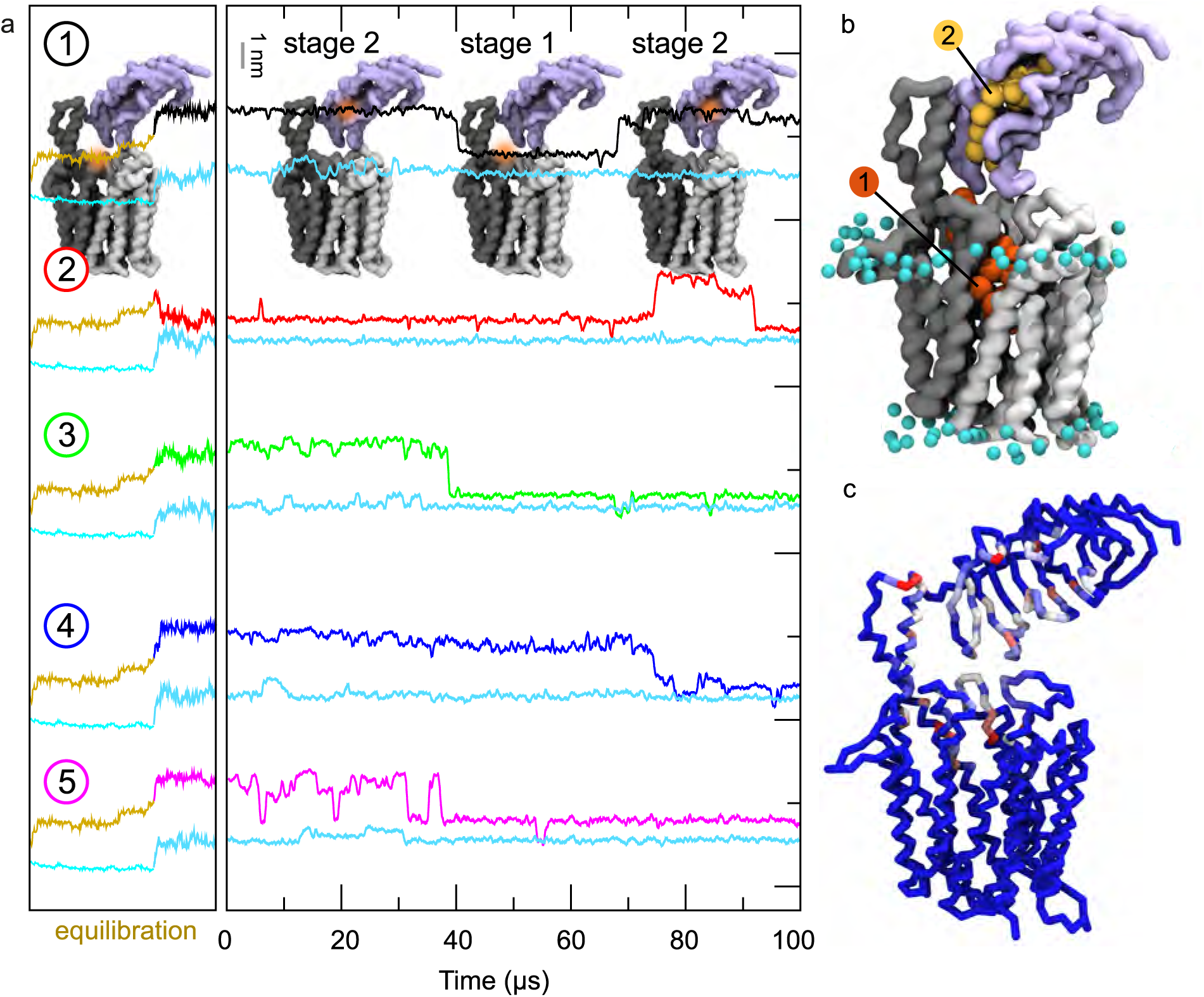
TAG loading into LprG in the MD simulations of the MHAS 216S^OUT^-LprG complex. (a) For each of the five simulations (numbered 1-5), the upper line depicts the z-coordinate of the center of mass of the TAG molecule. The cyan lines below correspond to the z-coordinate of the average center of mass of the phosphate groups of the upper membrane leaflet. The gold line is the equilibration phase (see methods). The models of the MHAS2l68^OUT^-LprG complex shown in the top panel indicate the position of the TAG (orange shade). (b) Visualization of TAG loading from the transporter’s main cavity (stage 1) along the TM11-TM12 extension into the LprG hydrophobic cavity (stage 2). Note that the overlay of structures shown displays the position of one single TAG molecule at two different points in time during the MD simulation. (c) TAG contacts with MHAS2168^OUT^-LprG, coloured from blue (no contacts) to red (large number of contacts).

## Discussion

In this work, we present the first high resolution structure of a mycobacterial MFS transporter, enabling the distinction of unusual features relevant to its function. Rv1410 acts in concert with periplasmic lipoprotein LprG to export TAGs from the inner membrane towards the mycomembrane. In our MD simulations, TAGs, lacking a polar head group, did not form an orderly part of the two leaflets of the cytoplasmic membrane, but instead accumulated in the hydrophobic core of the lipid bilayer. From a biophysical point of view, the main energy barriers TAG faces during its journey from the core of the inner membrane to the mycomembrane are the crossing of the charged outer leaflet of the inner membrane and the polar environment of the periplasm.

Our proposed model of transport (Fig. 7) accounts for this in that Rv1410 and LprG provide a continuous series of hydrophobic cavities and surfaces to shield TAG from the bulk water and thus allow for facilitated passage from the inner membrane core to the lipid binding cavity of LprG. Our functional data suggest that the TAG molecule enters Rv1410 in its IF conformation from the cytoplasmic membrane either via the TM5-TM8 or the TM2-TM11 lateral opening (step 1 in Fig. 7). According to our inward-facing homology model, the cavity of Rv1410 is hydrophobic enough to accommodate TAG and other lipids. Once TAG enters the cavity, Rv1410 transits from IF to OF conformation (step 2 in Fig. 7). Whether this transition is driven by TAG binding or protonation/deprotonation events, is not deducible from our data. As an effect, the TAG binding cavity is remodelled such that it opens to the periplasm and the E147-R417 ion lock is formed, which is a unique hallmark and essential functional element of Rv1410 and its mycobacterial homologues. The constriction pushes the TAG away from the membrane core to the level of the outer leaflet, while the hydrophobic cavity shields it from the charged environment of the lipid head groups. Thus, the TAG is enclosed in the transporter and retained in the cavity in the OF conformation because the lateral cleft between TM5-TM8 is narrow (Extended Data Fig. 2b) and the TM2-TM11 crevice is blocked by linker helices A and B. In fact, the discovery that linker helices of Rv1410 are crucial for its function is the first of its kind among 14-helix MFS transporters.

**Figure 7.**
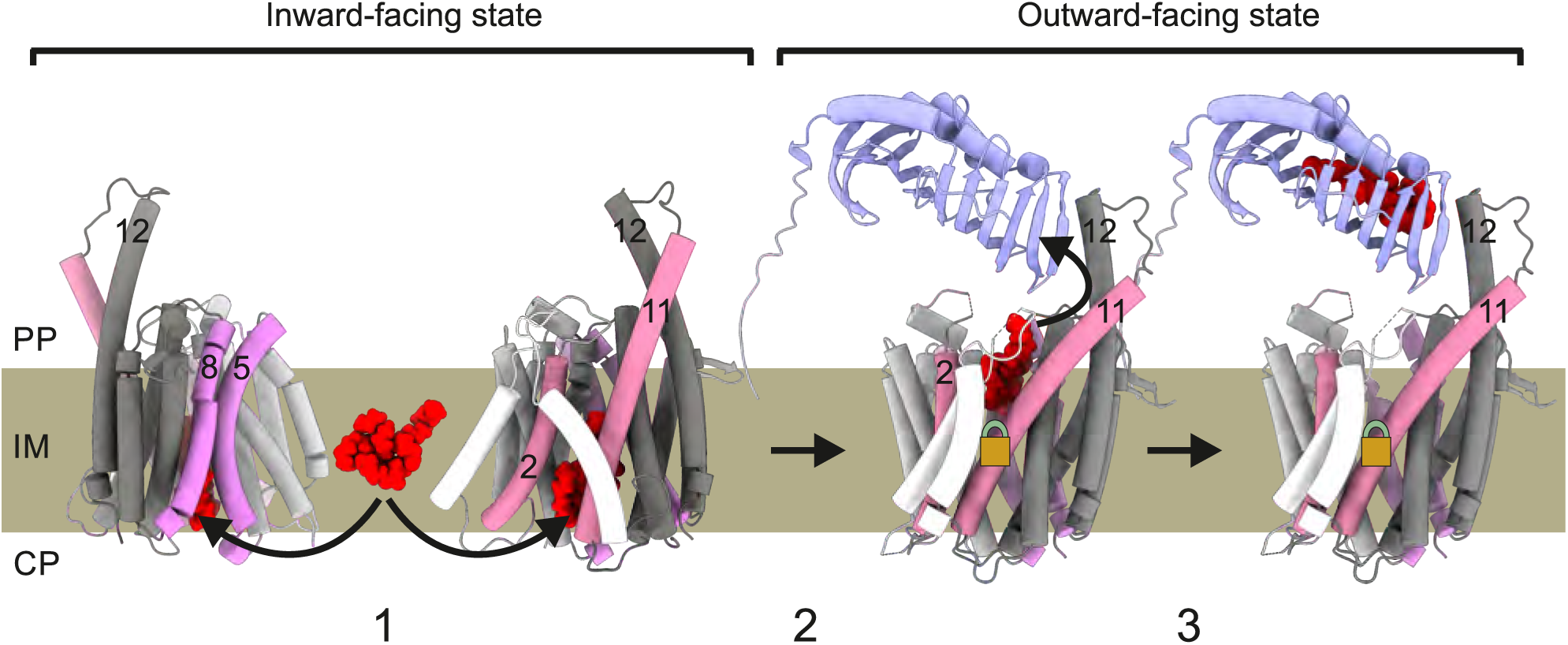
Mechanism of TAG transport by Rv1410 and LprG. Rv1410/MHAS2168 color scheme the same as in Fig lb, but transmembrane helices 5 and 8 are colored purple and transmembrane helices 2 and 11are colored pink. LprG crystal structure (PDB ID: 4ZRA) is colored pale lilac. (1) TAG molecule (red) enters the transporter’s central cavity through lateral openings between TM5-TM8 and TM2-TM11 in the inward-facing conformation. (2) The transporter switches from inward-facing to outward-facing state while the TAG molecule is occluded within the central cavity. The E147-R417 ion lock (symbolized by golden lock) at the bottom of the central cavity is formed and lifts TAG toward the periplasmic leaflet. (3) TM**11** and TM12 periplasmic extensions shield the TAG molecule from hydrophilic periplasm while the TAG molecule relocates from the transporter’s cavity tothe hydrophobic pocket of LprG.

As the last step of the transport pathway (step 3 in Fig. 7), TAG leaves the central cavity via the opening to the periplasm. The TAG remains concealed from the hydrophilic environment from one side by the TM11 and TM12 periplasmic extensions and from the other side by LprG which captures the lipid into its hydrophobic pocket. This final TAG extraction step is experimentally supported by the fact that TM12 truncations result in complete loss of transport activity, without disturbing transporter folding and production. Once TAG has left Rv1410, the proton gradient could be exploited to revert to the IF conformation and a new transport cycle can begin.

We discovered two functionally important carboxylate-arginine pairs in Rv1410, namely D22-R108 placed within the N-domain and E147-R417 placed across the N- and C-domain in the central cavity, whose most likely function is to couple proton influx to the export of TAG. It should be noted that the substitutions of the key carboxylates by the respective carboxamides did not fully abrogate transport function in MHAS2168 as it did in Rv1410 (Fig. 2c,d). This might be explained by the fact that TAGs are transported along their concentration gradient, as they are produced within the cell and finally dilute out to the mycomembrane and, to some extent, the growth medium. Hence, the transporter might well be able to facilitate TAG export without proton coupling, as has been shown in the case of several MFS transporters and their substrates^39,44^. However, the transport rate likely increases when lipid export is coupled to the proton gradient. The fact that the R426A_MH_ mutation is more deleterious than E157Q_MH_, might be due to an absolute requirement for the structural interaction at the base of the OF cavity between the two residues. Q157_MH_ is still able to form hydrogen bonds with R426_MH_, which is not possible for A426_MH_ with E157_MH_.

Extraction of lipopolysaccherides or lipoproteins from the outer leaflet of the inner membrane of gram-negative bacteria requires an active transport step mediated by the essential ABC transporters LptB_2_FG and LolCDE, respectively^45–47^. These lipid extractors belong to the type VI ABC transporters^48^, and utilize the energy of ATP binding and hydrolysis to pass their hydrophobic cargo to the dedicated periplasmic carrier protein. In this work, we describe the first structure of an MFS transporter capable of lipid extraction, which uses the proton-motive force instead of ATP to energize transport.

In contrast to LptB_2_FG and LolCDE which both directly interact with their cognate periplasmic proteins LptC^45^ and LolA^49^, respectively, LprG and Rv1410 do not appear to engage in strong and specific protein-protein interactions, as shown by the functional operon shuffling experiments (Fig. 5d-h) and previously described biochemical studies^8^. We therefore surmise an interplay in which LprG scans the periplasmic surface of the inner membrane while being attached via its lipid anchor until it encounters Rv1410 presenting a TAG molecule, ready to be captured into the lipid binding cavity of LprG. Whether LprG passes TAG onto further periplasmic proteins or whether it can be extracted itself from the inner membrane to channel TAG to the mycomembrane is currently unclear and requires further investigation.

## Methods

### Bacterial strains, media and plasmids

In this study, *Escherichia coli* strains DB3.1 and MC1061 were used for cloning and Rv1410, nanobody, and megabody expression, *M. smegmatis* MC^2^ 155 harboring pACE_C3GH_MHAS2168 was used for MHAS2168 protein expression and *M. smegmatis* MC^2^ 155 Δ*MSMEG3069/70* (dKO) was used for complementation studies. Mycobacteria were grown at 37°C in liquid Middlebrook 7H9 medium containing 0.05% Tween 80 supplemented with OADC or on solid Middlebrook 7H10 medium supplemented with OADC containing 4.5 ml/l glycerin. For MHAS2168 expression, 7H9 medium supplemented with 0.05% Tween 80 and 0.2% glycerol was used. *E. coli* was grown in lysogeny broth medium (LB) or Terrific broth medium (TB) at 37°C or 25°C respectively. Where required, the liquid medium was supplemented with the following amounts of antibiotics: 100 μg/ml of ampicillin (Amp^100^) and 25 μg/ml of chloramphenicol (Cm^25^) for *E. coli*, 50 μg/ml apramycin (Apr^50^) for *E. coli* and *M. smegmatis*, 50 μg/ml hygromycin B (Hyg^50^) for *E. coli* and *M. smegmatis*. Solid LB medium was supplemented with 120 μg/ml of ampicillin (Amp^120^), 20 μg/ml of chloramphenicol (Cm^20^), 50 μg/ml apramycin (Apr^50^) or 100 μg/ml hygromycin B (Hyg^100^). 7H10 medium was supplemented with 50 μg/ml apramycin (Apr^50^) or 50 μg/ml hygromycin B (Hyg^50^).

### Construction of plasmids

#### Shuffled *lprG-mfs* plasmids

The pFLAG plasmids harboring wild-type *lprG-mfs* operons and its single genes from *M. tuberculosis*, *M. smegmatis*, and *M. abscessus* originated from our previous study^8^. The same FX cloning strategy^35^ was at first used for *M. hassiacum lprG-mfs* operon *MHAS2167/68* and its single genes, using primers from Table S5 and *M. hassiacum* strain DSM 44199 as a template for colony-PCR with Q5 High-Fidelity DNA polymerase (NEB) to generate a fragment cloned into initial vector pINIT (Cm^25^). However, since the *MHAS2167* gene contains a SapI cleavage site with identical overhang (AGT), it was inefficient to use FX cloning to transfer the gene products to pFLAG vector. Thus, pFLAG plasmids containing *MHAS2167* and *MHAS2167-rv1410c* operon were generated using CPEC^50^ (primers in Table S5).

Using the pINIT plasmids with wild-type *lprG-mfs* operons as templates, shuffled operons were assembled (with primers in Table S5) using CPEC protocol^50^. The new operon sequences were confirmed by Sanger sequencing and then the shuffled operons were transferred to pFLAG vector, using FX cloning^35^.

#### Rv1410 and MHAS2168 mutants in pFLAG vector

First, QuikChange site-directed mutagenesis protocol was used to introduce mutations into Rv1410 in pFLAG_Rv1410/11 or in MHAS2168 in pFLAG_MHAS2167/68, using primers (named mutation_FOR and mutation_REV) in Table S6. However, in many cases, it was difficult to obtain the necessary PCR products, presumably due to the length of the vector (∼6100-6300 bp) and GC content of the operons (64-65%). Therefore, CPEC protocol^50^ was used to assemble the constructs from two fragments. In each case, a larger “backbone” fragment was amplified using Q5 High-Fidelity DNA polymerase (NEB) with primers pFLAG_REV2 and mutation_FOR and a smaller “insert” fragment was amplified with primers pFLAG_FOR2 and mutation_REV (Table S6). Both QuikChange products and CPEC reaction products were transformed into *E. coli* MC1061, plasmids were extracted using a QIAprep Spin Miniprep Kit (QIAGEN) and the correct sequences were confirmed by Sanger sequencing.

#### Expression plasmids

The *MHAS2168* gene was transferred from pINIT_*MHAS2168* to the pACE_C3GH^35^ and *rv1410c* gene from pINIT_*rv1410c* to the pBXC3GH^51^ using FX cloning, resulting in pACE_C3GH_*MHAS2168* and pBXC3GH_*rv1410c* in which the *mfs* transporters are C-terminally fused to a 3C protease cleavage site, GFP and a His_10_-tag. The gene encoding nanobody H2 (Nb_H2) was transferred from pSb_init_Nb_H2 to pBXNPHM3^52^, resulting in pBXNPHM3_Nb_H2 in which the nanobody’s N-terminus is fused to the PelB signal peptide, His_10_-tag, maltose binding protein and a 3C protease cleavage site. To turn nanobody Nb_H2 into a megabody MB_H2 and Nb_F7 into Mb_F7, the genes encoding Nb_H2 and Nb_F7 and lacking the first thirteen N-terminal residues were transferred to pBXMBQ vector (a kind gift from Eric Geertsma and Benedikt Kuhn), using FX cloning. The resulting constructs pBXMBQ_MB_H2 and pBXMBQ_MB_F7 encode a fusion protein that consists of an N-terminal DsbA signal peptide, the first thirteen N-terminal residues that form the β-strand A of a nanobody, a scaffold protein (HopQ adhesin domain)^23^ and the rest of the nanobody residues (containing all three CDRs), fused C-terminally to a 3C protease cleavage site, His_10_-tag, and Myc-tag.

### Vancomycin sensitivity assays in complemented *M. smegmatis* dKO cells

High-throughput cellular growth assays to assess the functionality of the transporter and LprG were conducted in principle as described in our previous publication.^8^ In short, each tested strain was grown into stationary phase and diluted to OD_600_=0.4 in 7H9 medium. 10 µl of these diluted cultures were transferred to wells containing 1 ml 7H9 medium complemented with vancomycin (concentrations as described in main text and figures) in 4 technical replicates and the cultures were incubated at 37°C, 300 rpm in a 96-well plate. At indicated time-points, 50 µl of culture were removed from the growth plate, transferred to a microtiter plate and OD_600_ was measured in a PowerWave XS Microplate Reader (BioTek). The growth curves in the figures are representative of at least three biological replicates and error bars denote the standard deviation of four technical replicates. All biological replicates of Rv1410 and MHAS2168 mutants’ growth curves are shown on Extended Data Fig. 8.

The vancomycin concentrations used in different assays were calibrated according to the experimental aims. 0.08 µg/ml vancomycin enables the growth of empty vector control and single protein (transporter or lipoprotein) complementation strains within the experimental time frame (∼50 h). 0.1 µg/ml vancomycin concentration prevents the growth of the empty vector control and inactivated mutant, but allows separation of partially active mutants, such as previously characterized LprG V91W^12,13^ and Rv1410 Truncation 1(= Long loop)^8^. 0.4 µg/ml vancomycin was used in cases where we aimed at distinguishing Rv1410 mutants that display such small growth disadvantages compared to the wild type strain that no growth differences can be observed at milder 0.1 µg/ml vancomycin condition.

### Western blotting

Western blotting for FLAG-tag detection was carried out exactly as described previously.^8^

### Multiple sequence alignments

CLC Main Workbench was used to produce a multiple sequence alignment of Rv1410 homologues from 17 *Mycobacterium* species belonging to three phylogenetically separate clades: *M. abscessus*, *M. aurum*, *M. avium*, *M. chelonae*, *M. fortuitum*, *M. haemophilum*, *M. hassiacum*, *M. intracellulare*, *M. kansasii*, *M. leprae*, *M. marinum*, *M. phlei*, *M. saopaulense*, *M. smegmatis*, *M. tuberculosis*, *M. thermoresistibile*, *M. vaccae*. The alignment was visualized using JalView^53^. To prepare a multiple sequence alignment of different MFS transporters to test uniqueness of Rv1410/MHAS2168 features, MUSCLE algorithm^54^ was used for alignment of protein sequences acquired from PDB database and JalView was used for visualization of the multiple sequence alignment. The alignment was validated using a subset of the MFS transporters and their structure models by superimposition in Chimera.

### Structure predictions of Rv1410c and its homologues

ColabFold software^29^ employing AlphaFold2^55^ and MMseqs2^56^ was used in batch mode with default settings to predict the structures of Rv1410 homologues from the following mycobacterial species: *M. abscessus*, *M. aurum*, *M. avium*, *M. fortuitum*, *M. hassiacum*, *M. marinum*, *M. phlei*, *M. smegmatis*, *M. tuberculosis*, *M. thermoresistibile*. ColabFold was similarly used to predict the structures of Rv1410 and MHAS2168 helix truncation mutants. Best models in outward-open conformation were chosen, superimposed on each other, and hydrophobicity analysis was performed in UCSF Chimera^57^ and UCSF ChimeraX^58^.

### Nanobody selections

For the selection of Rv1410 or MHAS2168 specific nanobodies, an alpaca was immunized with subcutaneous injections four times in two-week intervals, each time with 200 µg purified Rv1410 or MHAS2168 in 20 mM Tris-HCl pH 7.5, 150 mM NaCl, and 0.03% (w/v) *n*-dodecyl-*β*-D-maltopyranoside (β-DDM). Immunizations of alpacas were approved by the Cantonal Veterinary Office in Zurich, Switzerland (animal experiment licence nr. 172/2014). Blood was collected two weeks after the last injection for the preparation of the lymphocyte RNA, which was then used to generate cDNA by RT-PCR to amplify the VHH/nanobody repertoire. Phage libraries were generated and two rounds of phage display were performed against transporters solubilized in β-DDM. After the final phage display selection round, 1023-fold enrichment was determined by qPCR using AcrB as background for MHAS2168 and 652-fold enrichment for Rv1410. The enriched nanobody libraries were subcloned into pSb_init^52^ by FX cloning and 95 single clones were analyzed per transporter by ELISA. In case of MHAS2168, out of 88 positive ELISA hits, 22 were Sanger sequenced and 14 unique nanobodies were chosen for purification and further analysis. In case of Rv1410, out of 44 positive ELISA hits, 24 were Sanger sequenced and 7 unique nanobodies were discovered.

### Expression and purification of *M. tuberculosis* Rv1410

Rv1410 was produced in and purified from *E. coli* MC1061 following the same protocol as described previously.^8^

### Expression and purification of *M. hassiacum* MHAS2168

*M. smegmatis* MC^2^ 155 preculture harboring pACE_C3GH_MHAS2168 was inoculated from glycerol stocks into 7H9, HygB^50^ and grown at 37°C for 4 nights. The preculture was diluted 1:25 (v/v) into fresh expression medium (7H9, 0.2% Glc, HygB^15^) and grown at 37°C until the culture OD_600_ reached 0.8-1.0 before induction of protein expression with 0.016% acetamide overnight. Cells were harvested for 20 min at 6,000 rpm in a F9-6×1000 LEX centrifuge rotor (ThermoScientific) at 4°C and resuspended in Resuspension Buffer (20 mM Tris/HCl pH 8.0, 200 mM NaCl) containing 3 mM MgSO_4_ and traces of DNaseI. Cells were snap-frozen in liquid N_2_ and stored at -80°C until membrane preparation. Membranes were prepared by homogenizing the cell suspensions with a pestle in a Dounce homogenizer to remove larger cell clumps and subsequently disrupting the cells with a Microfluidizer (Microfluidics) at 30 kpsi on ice. Unbroken cells and cell debris were removed by centrifugation for 30 min at 8,000 rpm in a Sorvall SLA-1500 rotor at 4°C. Membranes were collected in a Beckman Coulter ultracentrifuge using a Beckman Ti45 rotor at 38,000 rpm for 1 h at 4°C and resuspended in TBS (pH 7.5) containing 10% glycerol. Membranes were snap-frozen in liquid N_2_ and stored at -80°C until protein purification. Then, membranes were solubilized for 2 h using 1% β-DDM (w/v) and insolubilized material was removed by ultracentrifugation. The supernatant was loaded on Ni^2+^-NTA columns after addition of 15 mM imidazole, washed with Wash Buffer I (50 mM imidazole (pH 7.5), 200 mM NaCl, 10% glycerol, 0.03% (w/v) β-DDM) and eluted with Elution Buffer II (200 mM imidazole (pH 7.5), 200 mM NaCl, 10% glycerol, 0.03% (w/v) β-DDM). In order to remove the C-terminally attached GFP/His_10_-tag, the buffer of the protein preparation was first exchanged to SEC Buffer (20 mM Tris/HCl pH 7.4, 150 mM NaCl, 0.03% (w/v) β-DDM) via a PD-10 desalting column. In a second step, 3C protease cleavage was performed overnight. Finally, cleaved MHAS2168 was again loaded on a Ni^2+^-NTA column and washed out with SEC buffer to remove GFP/His_10_-tag and the His-tagged 3C protease. Then, it was either separated by size exclusion chromatography (SEC) on a Superose 6 Increase 10/300 GL column in SEC Buffer before Mb_H2 complex formation or added to nanobody H2 to form a complex.

### Expression and purification of Nb_H2

*E. coli* MC1061 preculture harboring nanobody H2 expression vector pBXNPHM3_MHAS_H2 was directly inoculated from glycerol stock into LB, Amp^100^ and grown at 37°C overnight. The preculture was diluted 1:40 (v/v) into fresh expression medium (TB, Amp^100^) and grown for 2 h at 37°C and an additional hour at 25°C before induction of protein expression with 0.02% L-arabinose overnight. Cells were harvested for 20 min at 6,000 rpm in a F9-6×1000 LEX centrifuge rotor (ThermoScientific) at 4°C and resuspended in Resuspension Buffer containing 3 mM MgSO_4_ and traces of DNaseI. Cells were disrupted with a Microfluidizer (Microfluidics) at 30 kpsi on ice and unbroken cells and cell debris were removed as described above. Imidazole, to a final concentration of 20 mM, was added to the supernatant which was loaded on Ni^2+^-NTA columns. The columns were washed with Wash Buffer II (1x TBS buffer (pH 7.5), 50 mM imidazole) and the bound nanobody was eluted with Elution Buffer II (1x TBS (pH 7.5), 300 mM imidazole). The eluted protein was dialyzed against SEC buffer overnight at 4°C to remove excess imidazole and simultaneously cleaved with 3C protease to remove the N-terminally fused maltose binding protein and His-tag. Finally, nanobody H2 was again loaded on a Ni^2+^-NTA column and washed out with 1x TBS (pH 7.5) containing 40 mM imidazole to remove MBP and His10-tag and the His-tagged 3C protease. The protein was then snap-frozen in liquid N_2_ and stored at -80°C until it was separated by SEC (Sepax SRT-10C-300) in SEC buffer.

### Expression and purification of MB_H2 and MB_F7

*E. coli* MC1061 preculture harboring megabody expression vector pBXMBQ_MHAS_H2 or pBXMBQ_Rv_F7 was directly inoculated from glycerol stock into LB, Amp^100^ and grown at 37°C overnight. Megabody F7 expression and purification were conducted as described for nanobody H2 with the exception of the last SEC step when it was separated on a Superdex 200 10/300 GL column. Megabody H2 expression, cell harvest, cell disruption and first purification steps were carried out as described for nanobody H2. After first elution in Elution Buffer II, the megabody sample was contaminated with DNA, therefore the sample was dialyzed against 1x TBS (pH 7.5) overnight, then 3 mM MgSO_4_ was added and DNase treatment was performed for 2 h at 4°C. Again, the sample was loaded on Ni^2+^-NTA columns which were washed with Wash Buffer II and the bound megabody was eluted with Elution Buffer II. To remove the C-terminally attached His_10_-tag, the buffer was exchanged to 1x TBS (pH 7.5) via a PD-10 desalting column and then 3C protease cleavage was performed overnight. Then, megabody H2 was again loaded on a Ni^2+^-NTA column and washed with 1x TBS (pH 7.5) containing 30 mM imidazole to remove the His10-tag and the His-tagged 3C protease. Finally, it was separated by SEC on a Superose 6 Increase 10/300 GL column in SEC Buffer.

### Crystallization of MHAS2168 & Nb_H2 complex

Rv1410 did not yield any crystals after extensive vapour diffusion crystallization screening. Therefore, we purified its homologues from thermophilic mycobacterial species *M. thermoresistibile* and *M. hassiacum* and attempted to crystallize them. MHAS2168, the *M. hassiacum* Rv1410 homologue, produced crystals diffracting up to 7 Å. Subsequently, we generated 14 nanobodies against MHAS2168 and with three of these nanobodies, we obtained crystals diffracting up to 4 Å. Finally, systematic crystallization screening of the three MHAS2168-nanobody complexes in lipidic cubic phase (LCP) produced several crystals of the MHAS2168-Nb_H2 complex diffracting up to 2.7 Å, resulting in two native datasets (Table S1). To obtain the MHAS2168-Nb_H2 complex in LCP, Nb_H2 was separated on a Sepax SRT-10C-300 column and mixed with MHAS2168 in a molar ratio of 1:1.5 (transporter:nanobody). After incubation on ice (10 min), the complex was separated by SEC on a Superdex 200 10/300 GL column. The monodisperse peak of the complex was collected and concentrated with a 50 kDa cut-off concentrator (Vivaspin 2, Sartorius) and subsequently used for crystallization in LCP.

The concentrated transporter-nanobody complex (35 mg/ml) was mixed with molten 1-Oleoyl-rac-glycerol (monoolein, Sigma-Aldrich) at a protein:lipid ratio of 2:3 (v/v) using coupled syringe devices. 37 nl LCP boli were dispensed with a Crystal Gryphon LCP (Art Robbins Instruments) onto 96-well glass bases with a 120 µm spacer (SWISSCI), overlaid with 800 nl precipitant solution and sealed with a cover glass. The crystals were grown at 20°C and reached full size by day 12. Two native datasets were obtained from 3 crystals (I, II, III) grown in different reservoir solutions: I – 360 mM (NH_4_)H_2_PO_4_, 0.1 M sodium citrate (pH 6.3), 31% (v/v) PEG400; II – 380 mM NaH_2_PO_4_, 0.1 M sodium citrate (pH 5.7), 28% (v/v) PEG400, 2.4% (v/v) 1,4-butanediole; III – 420 mM NaH_2_PO_4_, 0.1 M sodium citrate (pH 5.8), 28% (v/v) PEG400, 2.4% (v/v) 1,4-butanediole. X-ray diffraction data were collected at the X06SA beamline (Swiss Light Source, Paul Scherrer Institute, Switzerland) on an EIGER 16M detector (Dectris) with an exposure setting of 0.05 s and 0.1° of oscillation over 120°. Diffraction data was processed with the XDS program package^59^ and datasets from crystals II and III were merged with xscale from the XDS program package^59^. Crystal I produced a complete dataset with no need for merging. The data-processing statistics are summarized in Table S1. Both native datasets showed diffraction to 2.7 Å.

### Cryo-EM analysis of MHAS2168 & Mb_H2 complex

After separation of both the transporter and megabody alone on a Superose 6 Increase 10/300 GL column in SEC Buffer, monodisperse peaks of both proteins were gathered and mixed in molar ratio of 1:1.2 and incubated on ice for 10 minutes. After concentration with a 100 kDa cut-off concentrator (Amicon Ultra-0.5 Centrifugal Filter Unit) to remove empty β-DDM micelles, the MHAS2168-Mb_H2 complex was separated again on a Superose 6 Increase 10/300 GL column in SEC Buffer and a monodisperse peak was collected and concentrated for cryo-EM analysis. 4 µl of MHAS2168-Mb_H2 complex (9.4 mg/ml and 6 mg/ml) were applied to glow-discharged (45 s) holey carbon grids (Quantifoil R1.2/1.3 Au 200 mesh) and a Grid Plunger GP2 (Leica) was used to remove excess sample by blotting to filter paper (2.5-3.5 s, 90-95% humidity, 10°C) and to plunge-freeze the grid rapidly in liquid ethane. The grids were stored in liquid N_2_ for data collection. The samples were imaged on a Titan Krios G3i (300 kV, 100 mm objective aperture), using a Gatan BioQuantum Energy Filter with a K3 direct electron detection camera (6k x 4k pixels) in super-resolution mode. 11,713 micrographs were recorded with a defocus range of -1 to -2.5 µm in an automated mode using EPU 2.7. The dataset was acquired at a nominal magnification of 130,000x, corresponding to a pixel size of 0.325 Å per pixel in super-resolution mode, with the total accumulated exposure of 66.54 e^-^/Å^2^ fractionated into 37 frames.

The data was processed in cryoSPARC v3.2^60^. First, the micrographs were subjected to patch motion correction and Fourier cropping, resulting in a pixel size of 0.65 Å per pixel. After subsequent patch CTF estimation, 11,542 good quality micrographs were selected, based on estimated resolution of CTF fits, relative ice thickness, and total full-frame motion. Template picking was used to pick particles from the micrographs; the templates were produced from an earlier lower-resolution map of MHAS2168-Mb_H2 complex. Particles were extracted with a box size of 600 pixels and Fourier-cropped to 300 pixels. After 6 rounds of 2D classification, 733,891 particles were subjected to 3-class *ab initio* reconstruction (default parameters) and 546,068 particles from the best two classes, showing megabody binding to the transporter, were used as input for heterogeneous refinement with the best-resolved and worse-resolved classes used as references and other parameters set to default. 402,229 particles from the best-resolved class (FSC resolution 7.19 Å) were directed into non-uniform refinement^61^ which produced a map of the complex resolved to 4.2 Å, according to the 0.143 cut-off criterion^62^. To further improve the resolution, the particle set was extracted again from the micrographs with a 450-pixel box size without Fourier cropping and subjected to non-uniform refinement (default parameters). This resulted in a 4.0 Å cryo-EM map.

### Cryo-EM analysis of Rv1410 & Mb_F7 complex

Rv1410-Mb_F7 complex was prepared similarly to MHAS2168-Mb_H2 complex, but the sample concentration was 4.4 mg/ml and it was applied to holey carbon grids with copper mesh (Quantifoil R1.2/1.3 Cu 200 mesh).

The data was acquired and processed similarly to MHAS2168-Mb_H2 complex, with the exception of recording 7,984 micrographs with electron dose of 65.0 e^-^/Å^2^ fractionated into 48 frames or 55.0 e^-^/Å^2^ into 38 or 41 frames. After template picking of particles from 7,631 good quality micrographs and 5 rounds of 2D classification, 427,971 particles were subjected to 3-class *ab initio* reconstruction (default parameters) and 300,246 particles from the best two classes, showing megabody binding to the transporter, were used as input for heterogeneous refinement with the best-resolved and worse-resolved classes used as references and other parameters set to default. 127,196 particles from the best-resolved class (FSC resolution 8.64 Å) were directed into non-uniform refinement^61^ which produced a map of the Rv1410-Mb_F7 complex resolved to 7.51 Å, according to the 0.143 cut-off criterion^62^.

### Structure determination

SWISS-MODEL^63^ was used to generate homology models of MHAS2168 (based on PDB structure 6GS4) and Nb_H2 (based on PDB structure 5F7L) which were trimmed into polyalanine models with the CHAINSAW^64^ program from CCP4 suite^65^ and fitted into the 4.0 Å cryo-EM map in *Coot*^66^. ColabFold^29^ software, combining MMseqs2 with AlphaFold2^55^, was used to predict the structure of Rv1410 and several of its mycobacterial homologues. In *Coot*^66^, registry was established and side-chains built manually into well-resolved helices 1-12 in the model fitted into the 4.0 Å cryo-EM map, while comparing the map to ColabFold structure predictions. The model was subjected to one cycle of real-space refinement in Phenix^67^ before using it as a search model in molecular replacement with Phaser ^68^ to phase the native 2.7 Å single-crystal dataset. After iterative cycles of refinement and modelling with phenix.refine^69^ and ISOLDE^70^, a model was produced whose R-factors (R_work_=0.2708 and R_free_=0.3277) could not be improved with further refinement. The better-resolved chains A (MHAS2168) and B (Nb_H2) from this model were used as search model for molecular replacement with Phaser MR to phase the native 2.7 Å merged dataset. Again, refinement and modelling was performed with phenix.refine^69^ and ISOLDE^70^ to reach the final R-factors (R_work_=0.2450 and R_free_=0.2915) as indicated in Table S1. The structure was validated with Molprobity^71^.

### Fitting a TAG molecule into the 4.0 Å cryo-EM map

A TAG molecule, tripalmitoylglycerol (4RF) was fitted to the non-proteinaceous density in the 4.0 Å cryo-EM map by visual evaluation, using Coot and ISOLDE. First, chains A (MHAS2168) and B (Nb_H2) of the crystal structure were fitted into the 4.0 Å cryo-EM map in Chimera, then the TAG molecule was imported and fitted manually in Coot. Finally, polishing of the structure was performed in ISOLDE and final refinement in Phenix.

### Molecular dynamics simulations

Coarse-grained MD simulations of the MHAS2168 transporter from *Mycobacteria hassiacum* in outward-open conformation were carried out with GROMACS version 2021.1^72^ with the Martini 2.2 force field^73^. The sizes and compositions of all the simulated systems are listed in Table S3. From the X-ray crystal structure, missing residues 55-57, 203-208 and 234-236 were added to the chain A using MODELLER^74^, the best model was selected out of 100 structures. The model was briefly refined using ISOLDE^70^ and further oriented along the z-axis using the PPM web server^75^. The obtained MHAS2168 atomistic model described above was converted to the coarse-grained resolution using the martinize.py script. To maintain the structural integrity of the protein, an elastic network with a cutoff distance of 7 Å for the MHAS2168^OUT^, 9 Å for the MHAS2168^IN^, 9 Å for the MHAS2168^IN-TAG^ and 9 Å for the MHAS2168^OUT^-LprG (see below) was used with force constants of 1000 kJ × mol^-1^ × nm^-2^. For the MHAS2168^OUT^-LprG complex, 134 (out of a total of 3698) intra- and inter-molecular elastic network bonds were removed to avoid a too rigid protein-protein interface in the MHAS2168-LprG complex. The Martini Maker tool available in the CHARMM-GUI web server^76^ was used to insert the protein into a symmetric bilayer resembling the mycobacterial plasma membrane composition^3,4,77^ (see Table S3): 35% 1-palmitoyl-2-oleyl-phosphtidylethanolamine (POPE), 30% 1-palmitoyl-2-oleyl-phosphtidylglycerol (POPG), 15% cardiolipin (CDL), 10% 1-palmitoyl-2-oleyl-phosphtidylinositol (POPI) and 1% triacylglycerol (TAG)^78^. The MHAS2168^OUT^-LprG complex was embedded in a bilayer of equal composition but using the insane tool^79^, which was also used to build the Myco^mem^ system (see Table S3). For the MHAS2168^IN-TAG^ system, 1 TAG molecule was initially positioned within the main cavity of the transporter. The TAG molecule within the MHAS2168^OUT^-LprG complex was modeled with 2 hydrophobic tails pointing upwards (towards LprG) and, as a control simulation, only 1 tail pointing upwards and 2 tails downwards (towards MHAS2168). Prior to the MD simulations, to avoid a possible bias, one POPE and one POPG molecule were removed from the lower leaflet of the MHAS2016^IN^ system because they entered the main transmembrane cavity during the equilibration phase. All the systems were neutralized with a 150 mM concentration of NaCl and subsequently energy-minimized with steepest descent until machine precision. The minimized systems were equilibrated in two consecutive steps of 50 ns and 200 ns of MD simulation, first with all protein beads and second with the protein backbone beads restrained by harmonic potentials (force constants of 1000 kJ × mol^-1^ × nm^-2^). The time step for integrating the equations of motion in the coarse-grained simulations was 20 fs. The “new-RF” simulation parameters were used, as suggested by de Jong et al.^80^. Since the *M. hassiacum* is a thermophile^81^, the equilibrations and production simulations were performed at 330 K with protein, each lipid species and solvent separately coupled to an external bath using the v-rescale thermostat^82^ with coupling time constant τ_T_ = 1.0 ps. Pressure was maintained at 1 bar using the stochastic cell rescaling (c-rescale) barostat^83^ with semi-isotropic conditions (coupling time constant τ_p_ = 12.0 ps and compressibility 3.0 × 10^-4^ bar^-1^). For each system, five independent production simulations (each of length 100 microseconds) were generated using different random seeds for the initial velocities. For the Myco^mem^ system, the production run was 20 microseconds long. For the control simulations of the MHAS2168^OUT^-LprG complex, after the initial 100 microseconds of production run, the simulation was further extended by 180 microseconds, and 5 additional simulations of 50 microseconds were carried out with different random seeds for the initial velocities. Coordinates were saved to the disk every 400 ps. All analyses were carried out on the trajectories from the production runs. The GROMACS analysis tool gmx select in combination with an in-house script was used to calculate the protein-lipid contacts. A contact between a CG lipid headgroup bead and a protein bead was defined within a distance of 0.6 nm between the two. The tool gmx trajectory was used to extract the z-coordinate of the TAG center of mass and gmx density was used to calculate the density profile of the membrane components. The volume of the main transmembrane cavity was calculated using trj_cavity (https://sourceforge.net/projects/trjcavity/)^84^. For these calculations, we have used index files consisting of selections of protein residues encompassing the cavity: Residues 32-44, 60-75, 114-129, 153-164, 223-234, 288-303, 318-332, 388-403 and 420-438 were selected for the MHAS2016^OUT^ conformation. Residues 64-81, 122-137, 144-160, 288-300, 325-336, 395-408 and 414-431 were selected for the MHAS2016^IN^ conformation. Before the calculations, the five trajectories where concatenated and the proteins were superimposed over the backbone atoms of the residues listed above. Molecular graphics were generated with VMD 1.9.4 (http://www.ks.uiuc.edu/Research/vmd/)^85^. Data were plotted using Grace (http://plasma-gate.weiz-mann.ac.il/Grace/).

### Modelling the MHAS2168 in inward-open conformation

To build an inward open conformation of MHAS21668, MODELLER^74^ was used. Initially, a web frontend (http://www.ebi.ac.uk/Tools/msa/clustalo) to CLUSTAL Omega^86^ was used to align the target protein sequence of MHAS2168 to the proton-dependent oligopeptide transporter PepTSo2 from *Shewanella oneidensis*^87^ (sequence identity is 23%). The obtained alignment was manually curated to avoid fragmentation mostly in the region of the linker helices. The PepTSo2 X-ray structure PDB 4LEP (resolution 3.20 Å) was used as template in combination with an initial mock-model of MHAS2168 in inward-open conformation. To obtain this mock-model, the N- (residues 1-183) and C-terminal regions (residues 263-495) of MHAS2168 were independently superposed to the homologous parts of PepTSo2. During the modeling procedure the linker helices were restrained to assume a canonical α-helix conformation, however, their overall position differed from the template 4LEP. Therefore, to correctly position the target linker-helices, the best structural model out of 100 generated ones was further used for a second round of MODELLER together with both linker-helices modeled in the initial step and in turn superposed to the 4LEP linker-helices.

### Modelling the MHAS2168^OUT^-LprG complex

The LprG protein sequence from *M. hassiacum* was downloaded from Uniprot (id: K5BJY3) and the platform ColabFold^29^ was used to build the MHAS2168^OUT^-LprG heterodimeric complex. At variance of all the models obtained, the first best model features LprG quite far away from the extended helices TM11 and TM12, and consequently, the second best model was selected. Further, the refined atomistic MHAS2168 model described above was superposed over the TM11 and TM12 of the ColabFold MHAS2168 selected model and the first 30 N-terminal unstructured residues of LprG were removed.

## Acknowledgements

Dr. Simona Sorrentino of the Center for Microscopy and Image Analysis, University of Zurich, is acknowledged for help with cryo-EM grid preparation and cryo-EM data collection. Beat Blattmann and Caroline Müller of the Protein Crystallization Center, University of Zurich, are acknowledged for their help with setting up the crystallization screens. We thank Saša Štefanić for conducting alpaca immunizations and the staff of the SLS beamlines X06SA and X06DA for their support during data collection. We thank Jennifer C. Earp for help with cryo-EM grid preparation and Dr. Alisa Garaeva for her advice on cryo-EM data processing. We are very grateful to Dr. Eric Geertsma and Dr. Benedikt Kuhn for sharing the pBXMBQ vector with us. All members of the Seeger lab are acknowledged for project discussion. Work in the lab of MAS was supported by a SNSF Professorship of the Swiss National Science Foundation (PP00P3_144823), the European Research Council (ERC) (consolidator grant n° 772190) and a grant of the Novartis Foundation for Medical-Biological Research (to MAS). SR was supported by a Candoc fellowship of the University of Zurich (grant nr. FK-17-035). Work in the lab of LVS was supported by the Deutsche Forschungsgemeinschaft (DFG) under Germany’s Excellence Strategy – EXC 2033 – 390677874 – RESOLV and through grant SCHA1574/6-1.

## Author contributions

SR and MAS conceived the project. MH and SR cloned all genes into the respective complementation and expression vectors for *E. coli* and mycobacteria. MH screened Rv1410 in vapour diffusion experiments and initiated MHAS2168 screening. MH and SR purified protein for alpaca immunization. SR conducted alpaca nanobody selections. SR purified all proteins and protein complexes thereafter. SR and JS crystallized MHAS2168-Nb_H2 complex in LCP. SR, CAJH, IG, JS, and MAS collected crystal diffraction data. SR and CAJH processed the data and built and refined the model of MHAS2168-Nb_H2 complex. SR and IG prepared cryo-EM grids and performed cryo-EM data analysis. SR built and refined the MHAS2168 model into MHAS2168-Mb_H2 complex cryo-EM map. SR and MAS analyzed the structures and SR produced the multiple sequence alignments and structure predictions. SR designed and carried out the mutagenesis of Rv1410 and MHAS2168 and performed Western blotting. SR conducted vancomycin sensitivity assays. DD and LVS designed and performed the molecular dynamics simulations. DD generated the MHAS2168 inward-facing homology model and the MHAS2168-LprG complex. SN instructed SR in LCP methodology and contributed to crystallization strategy development. SR and DD prepared the figures and the tables. SR and MAS wrote the first draft of the manuscript. DD and LVS wrote the MD simulation sections. All authors edited the manuscript.

---

**Extended Data Figure 1.**
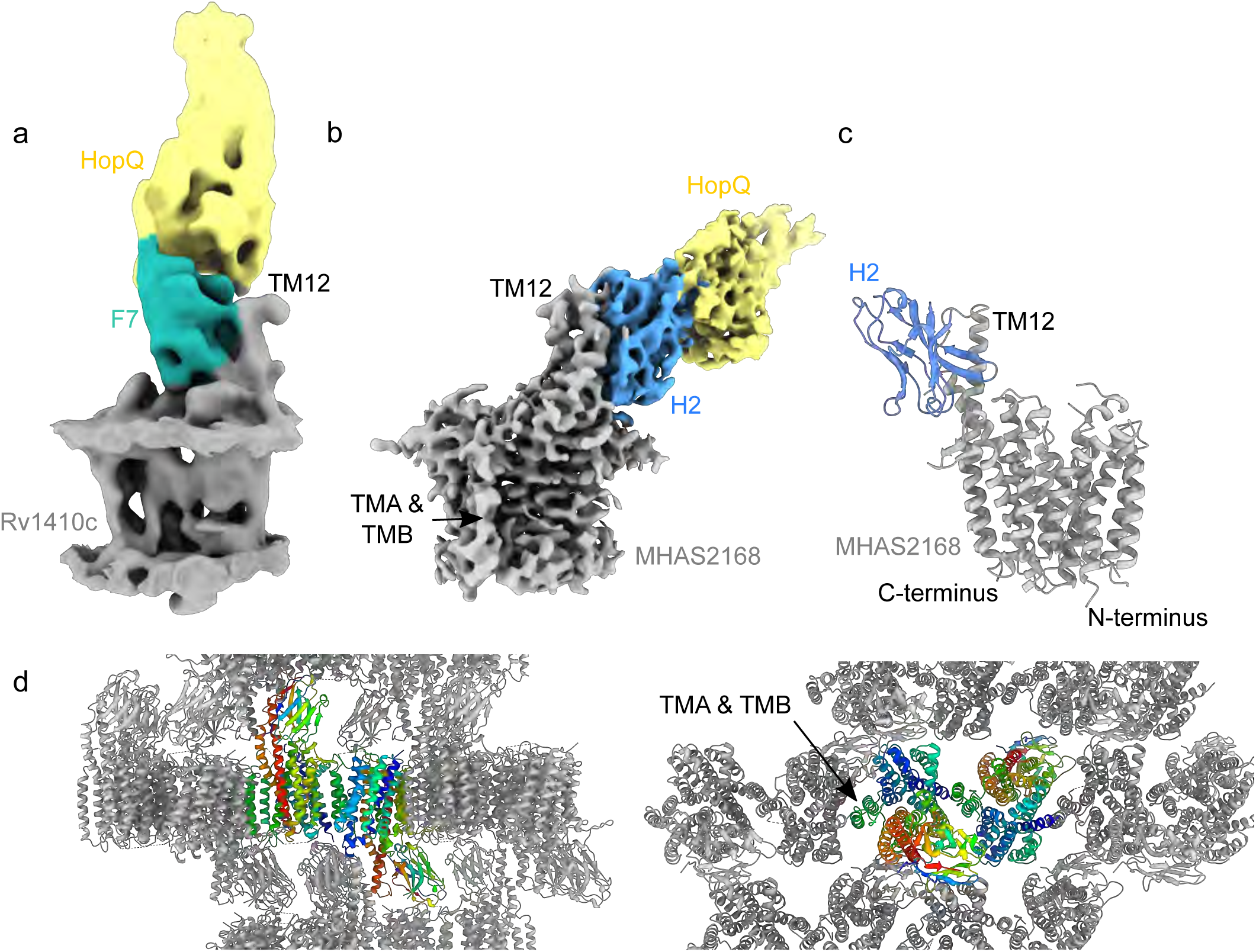
Determination of Rv1410 and MHAS2168 structures. (a) 7.5 Å cryo-EM map of Rv1410-Mb_F7 complex. Rv1410 – gray; Nanobody F7 – sea green; Megabody HopQ domain – yellow. (b) 4.0 Å cryo-EM map of MHAS2168-Mb_H2 complex. MHAS2168 – gray; Nanobody H2 – blue; Megabody HopQ domain – yellow. TMA & TMB – linker helices A and B. (c) 2.7 Å crystal structure of MHAS2168-Nb_H2 complex. MHAS2168 – gray; Nanobody H2 – blue. (d) Crystal packing of MHAS2168-Nb_H2 complex lipidic cubic phase (LCP) crystals. The asymmetric unit comprises two transporter/nanobody complexes (rainbow color scheme).

**Extended Data Figure 2.**
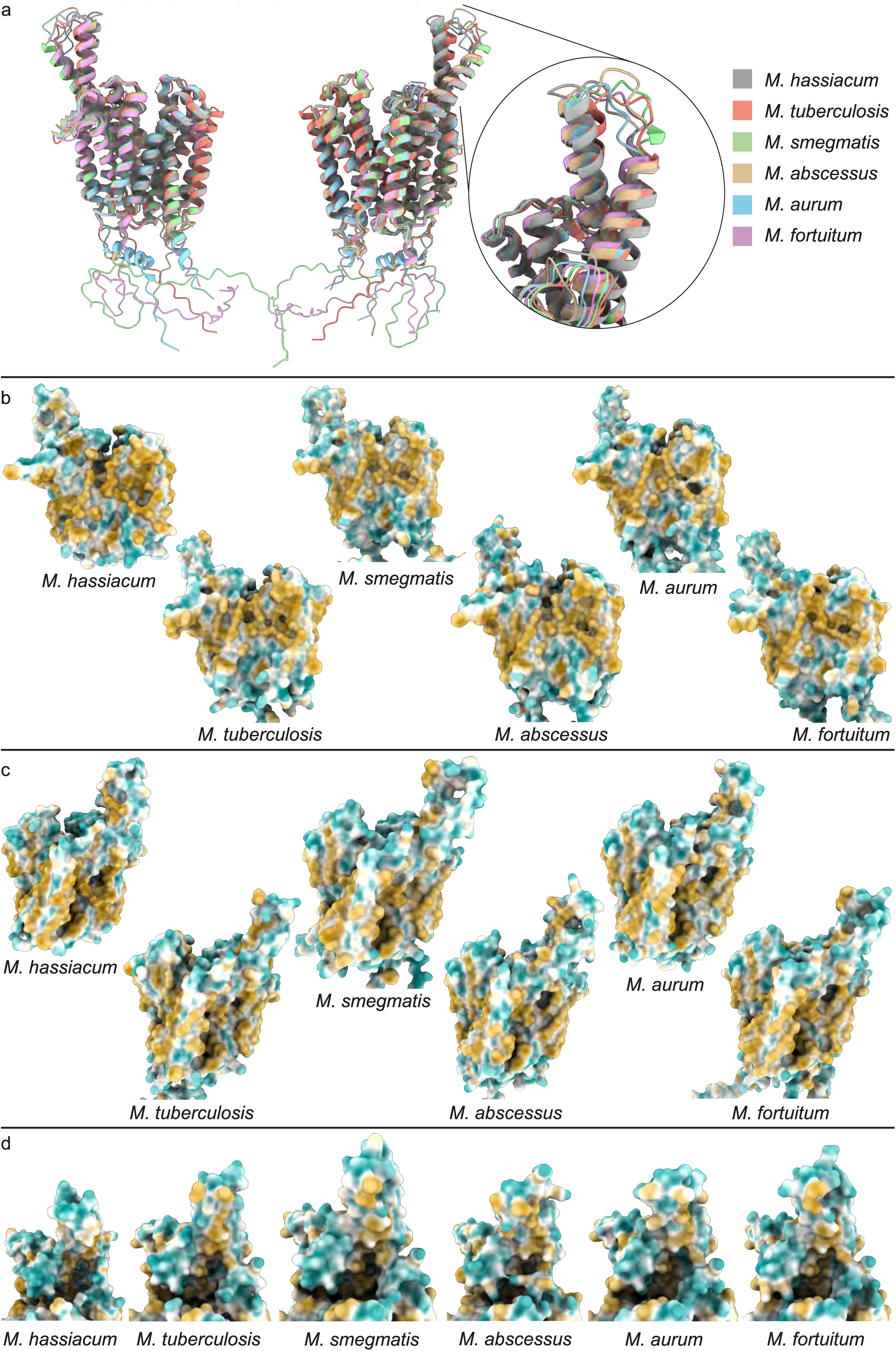
Comparison of MHAS2168 crystal structure and its mycobacterial homologues’ structure predictions. A subset of ColabFold structure predictions in outward-facing conformation that were used to analyze the common features of mycobacterial Rv1410/MHAS2168 homologues are shown together with the MHAS2168 crystal structure. (a) Structure predictions of homologues from *M. tuberculosis* (pale red), *M. smegmatis* (green), *M. abscessus* (beige), *M. aurum* (light blue), and *M. fortuitum* (pale purple) are superimposed on the crystal structure of MHAS2168 from *M. hassiacum* (gray). Inset: more detailed view of the TM11-TM12 periplasmic extensions shows differences in TM11 and loop length in different mycobacterial homologues while TM12 length remains the same. (b) Side views of hydrophobic surfaces of different mycobacterial homologues. The TM5-TM8 lateral openings are narrow and partially obstructed. (c) Opposite side views of hydrophobic surfaces of different mycobacterial homologues. Linker helices A and B are blocking the TM2-TM11 lateral opening. (d) Close-up view of the hydrophobic surfaces of the TM11-TM12 periplasmic extensions towards the central cavity. Hydrophobic patches are the common denominator of all periplasmic helix extensions. On panels (b) – (d), from left: homologues from *M. hassiacum*, *M. tuberculosis*, *M. smegmatis*, *M. abscessus*, *M.aurum*, *M. fortuitum*. On panels (b) – (d) hydrophobicity color scheme: hydrophobic – gold; hydrophilic – cyan.

**Extended Data Figure 3.**
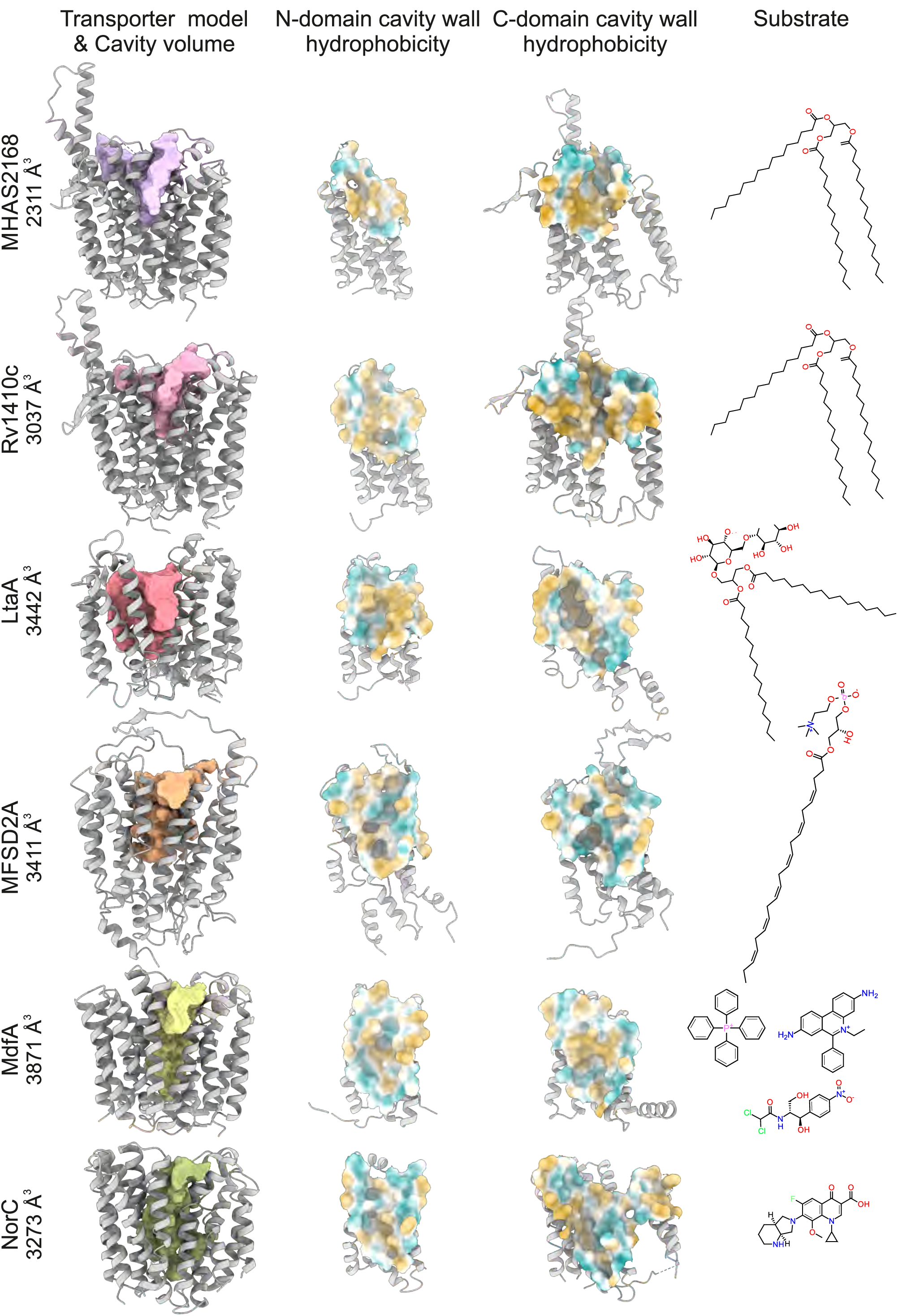
Comparison of central cavities from different MFS transporters known to transport lipids or drugs. The central cavity surface hydrophobicity/hydrophilicity reflects the polarity of the transporter’s substrate(s). Left column: Side views of transporters (gray) with their central cavity volumes highlighted in color. Middle left column: Inside view of the hydrophobicity surfaces of the central cavity walls from N-domains. Middle right column: Inside view of the hydrophobicity surfaces of the central cavity walls from C-domains. Right column: Structures of substrates transported by the corresponding MFS transporters. From top to bottom: TAG exporter MHAS2168 crystal structure (this study) and TAG species tripalmitoylglycerol; TAG exporter Rv1410 ColabFold structure prediction (this study) and TAG species tripalmitoylglycerol; Lipoteichoic acid lipid anchor flippase LtaA crystal structure (PDB ID: 6S7V); Lysophosphatidylcholine-docosahexaenoic acid importer MFSD2A cryo-EM structure (PDB ID: 7N98); Multi-drug efflux pump MdfA crystal structure (PDB ID: 6GV1) and its substrates tetraphenylphosphonium, ethidium, and chloramphenicol; Quinolone efflux pump NorC crystal structure (7D5P) and its substrate moxifloxacin. Hydrophobicity color scheme: hydrophobic – gold; hydrophilic – cyan.

**Extended Data Figure 4.**
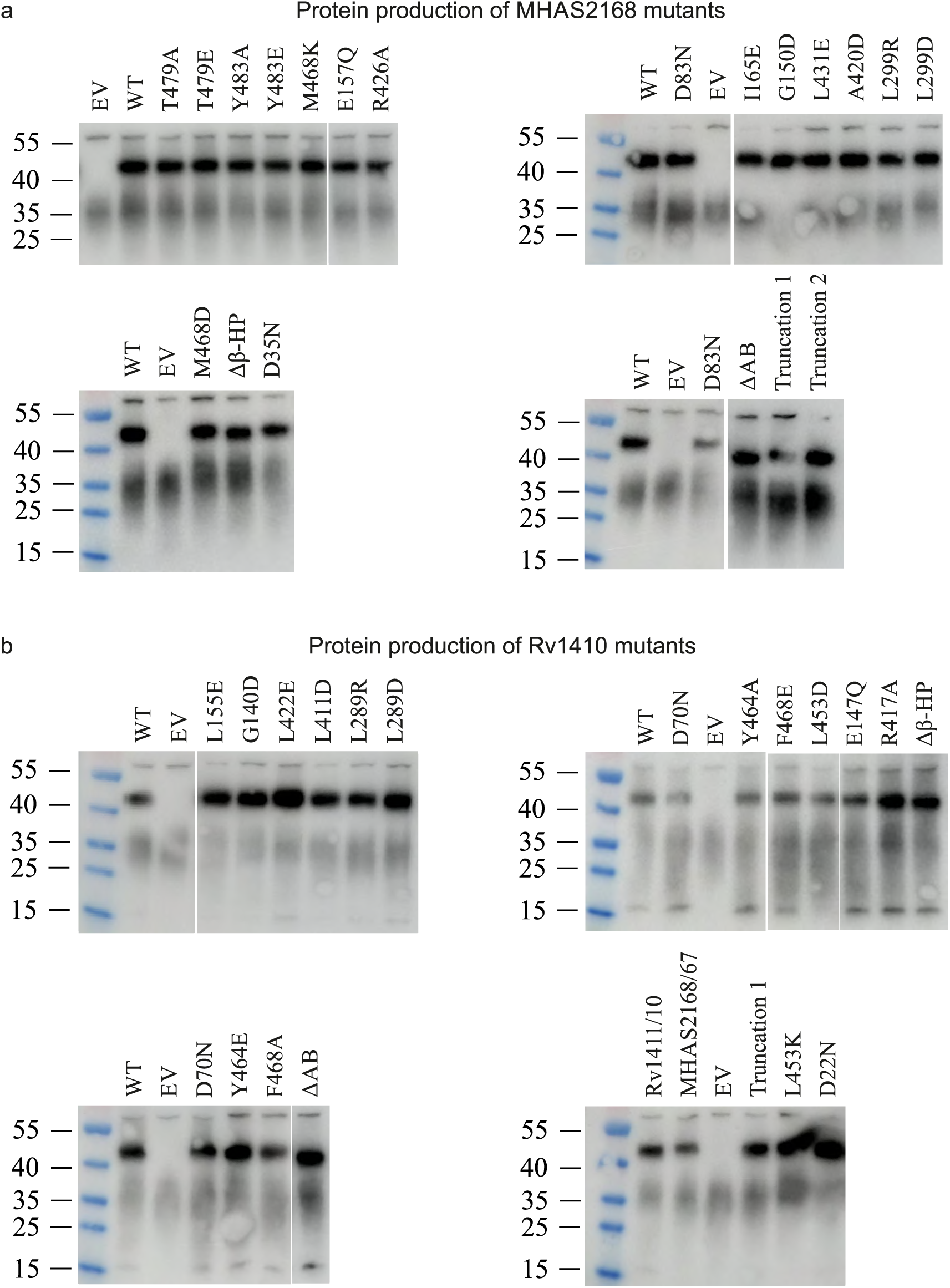
Protein production levels of Rv1410 and MHAS2168 mutants. Production levels of wild type MHAS2168, Rv1410, or their mutants expressed in *M. smegmatis* dKO using complementation vector pFLAG were probed by Western blotting via a C-terminal 3xFLAG tag. *M. smegmatis* dKO harboring the empty pFLAG vector served as negative control (EV). (a) Protein production levels of MHAS2168 and MHAS2168mutants.(b) Protein productionlevelsofRv1410andRv1410mutants.

**Extended Data Figure 5.**
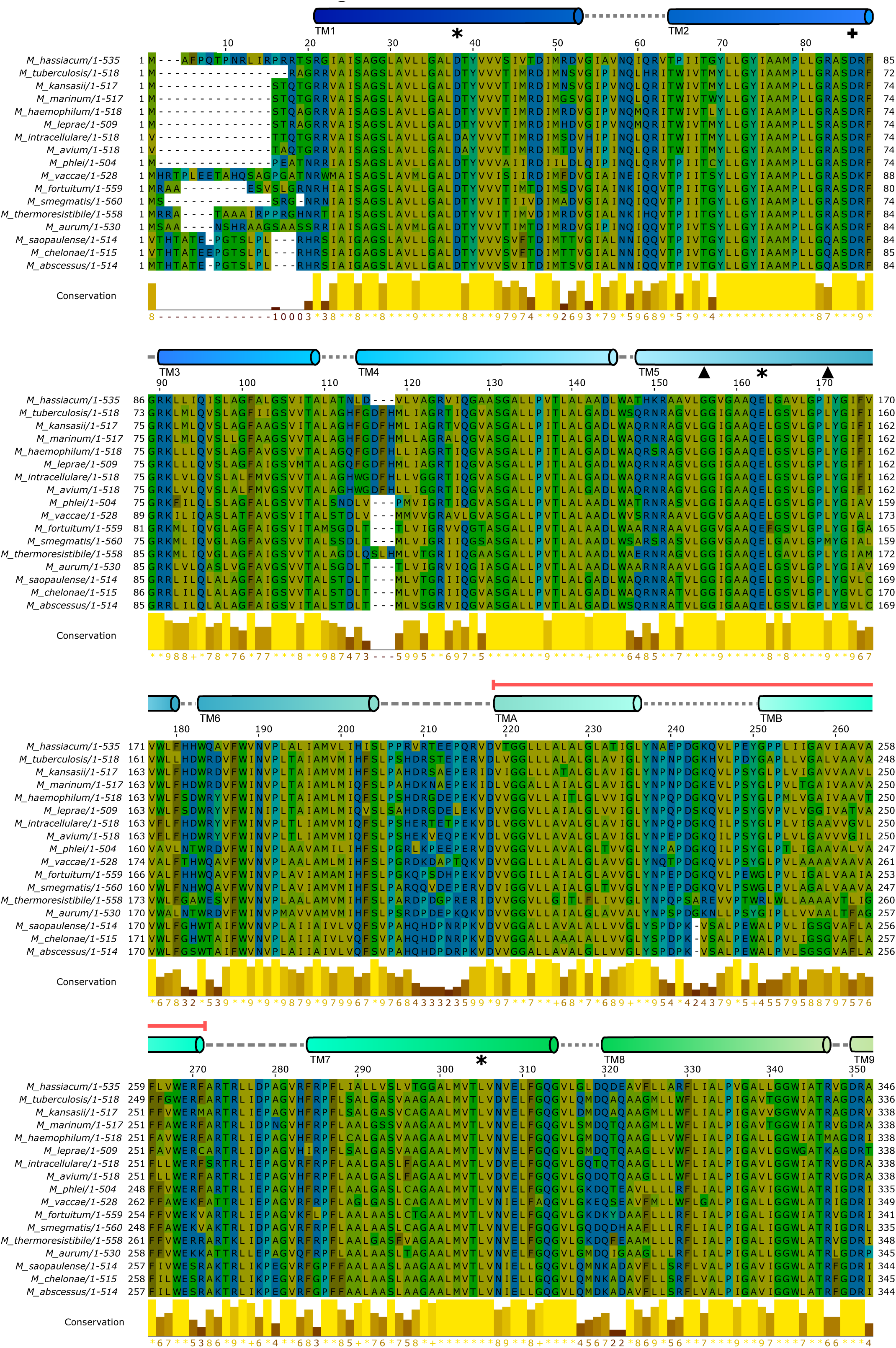
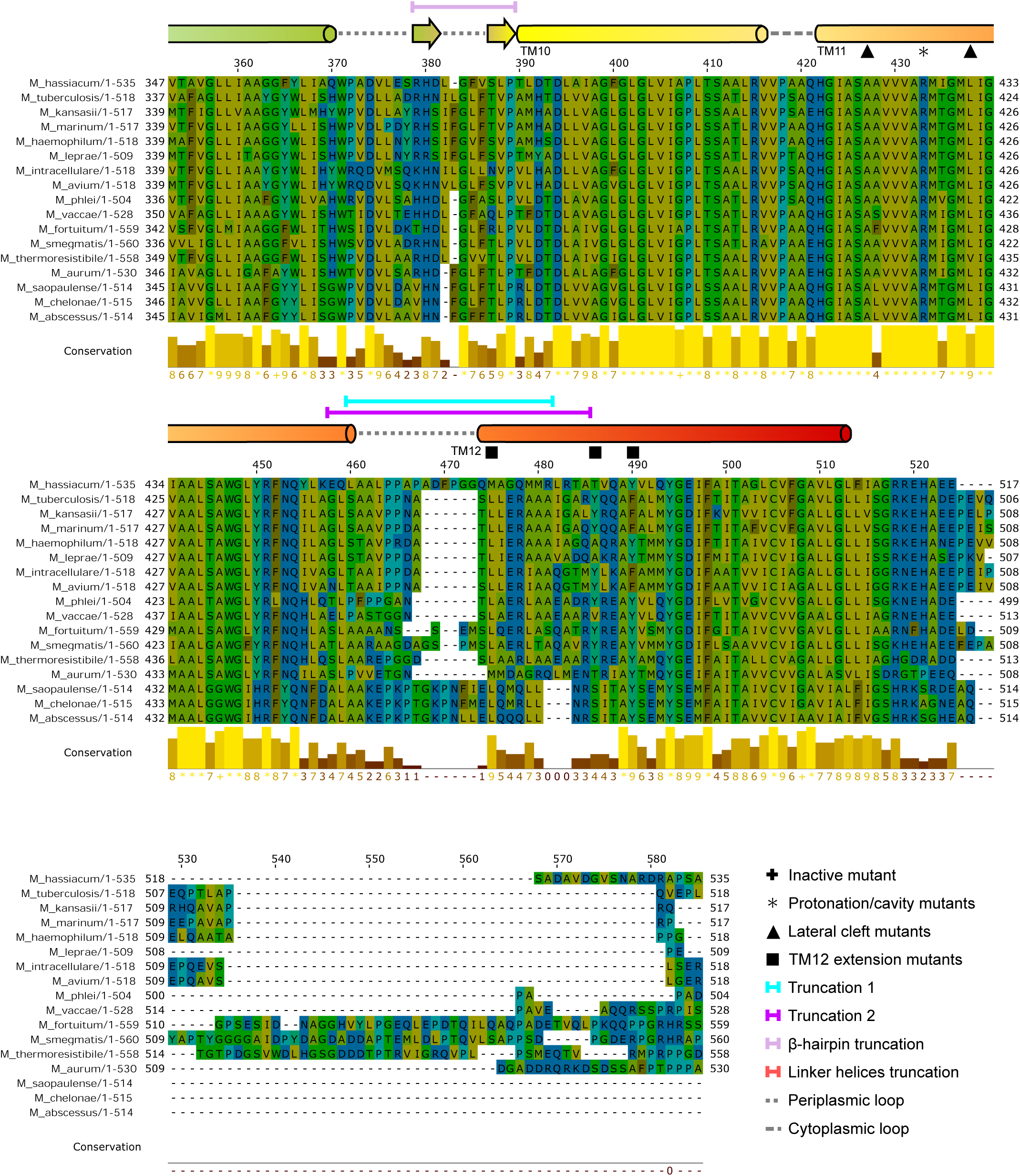
Multiple sequence alignment (MSA) of primary structures of 17 mycobacterial Rv1410 homologues. The amino acid residues in the MSA are colored according to their hydrophobicity (hydrophilic residues – blue; hydrophobic residues – gold). Secondary structure elements corresponding to the primary structure are depicted above the MSA: transmembrane α-helices and β-sheets are colored in the rainbow color scheme with the N-terminal end being blue and C-terminal end being red. Loops between secondary structure elements are depicted as gray dashed lines (periplasmic loops – short dashes; cytoplasmic loops – long dashes). Point mutations analyzed in this study are marked by different symbols above the corresponding amino acid residue in the MSA as indicated. Different truncation mutants are marked with colored lines showing the extent of truncations as indicated.

**Extended Data Figure 6.**
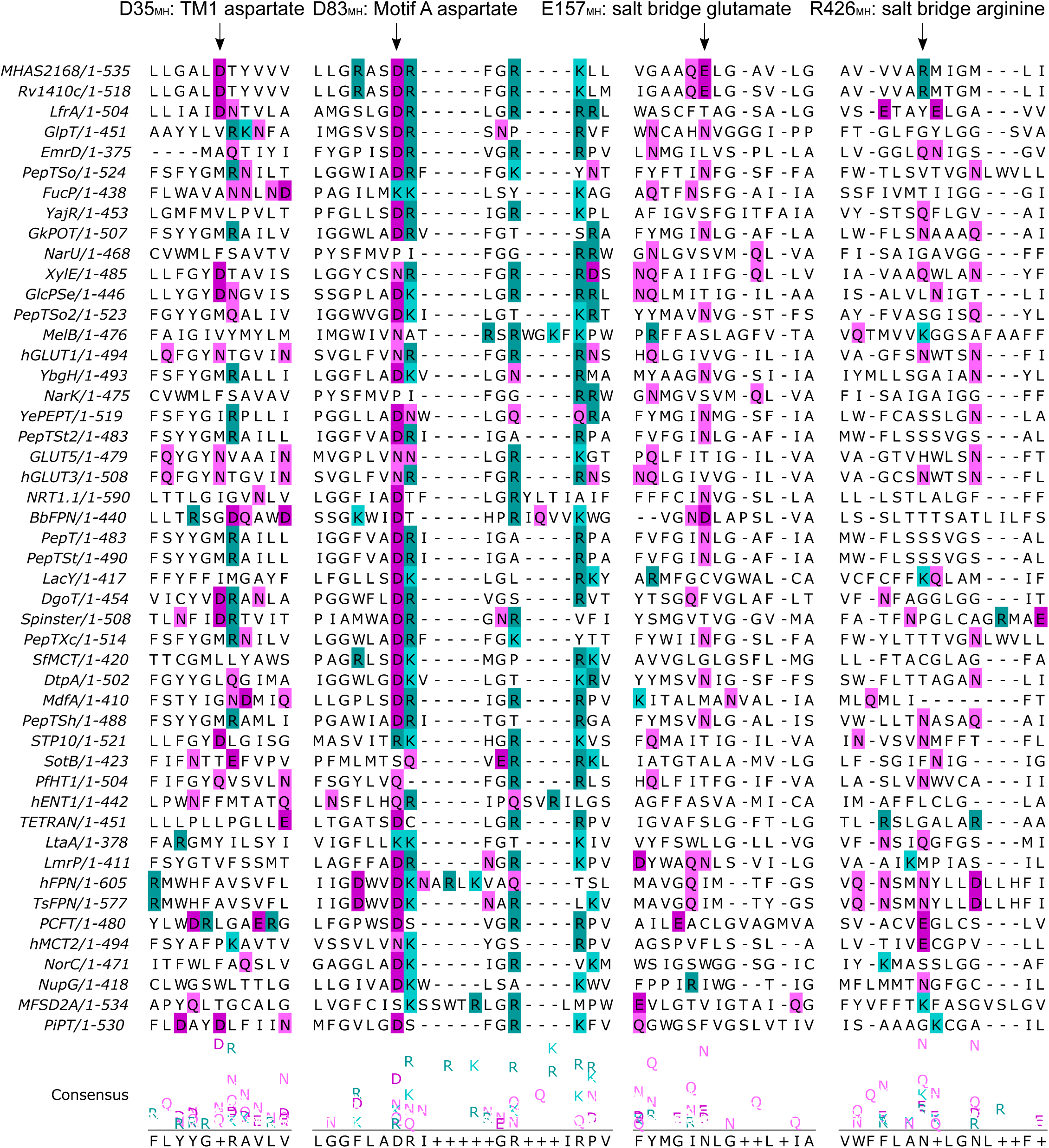
Conservation of different residues potentially involved in proton coupling in Rv1410 and MHAS2168 among other MFS transporters, depicted by a multiple sequence alignment. Positively charged residues arginine and lysine are highlighted in cyan (R and K). Negatively charged residues glutamate and aspartate are highlighted in dark pink (E and D). Glutamine and asparagine are highlighted in light pink (Q and N). 1st block: D35_MH_/D22_Mtb_ (TM1, side chain within the N-domain) is mostly conserved in mycobacterial MFS transporters and some sugar porters. 2nd block: Motif A aspartate D83_MH_/D70_Mtb_ (cytoplasmic loop between TM2 and TM3) is very conserved among MFS transporters. 3rd block: Salt bridge glutamate E157_MH_/E147_Mtb_ (TM5, side chain within the central cavity) seems to be present only in mycobacterial TAG exporters MHAS2168 and Rv1410. 4th block: Salt bridge arginine R426_MH/_R417_Mtb_ (TM11, side chain within the central cavity) seems to be present only in mycobacterial TAG exporters MHAS2168 and Rv1410.

**Extended Data Figure 7.**
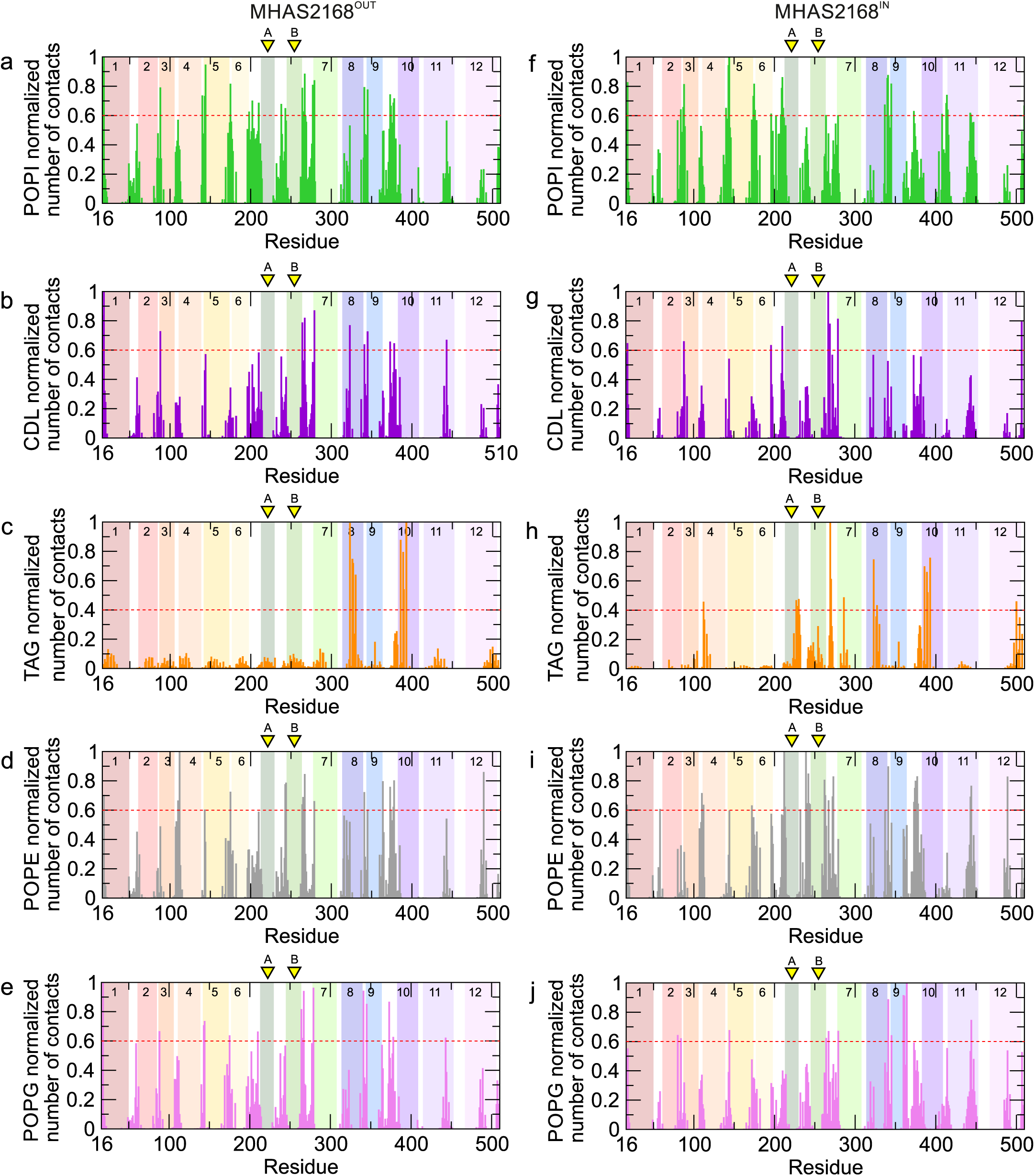
Lipid interactions observed in the simulations. Protein-lipid contacts were normalized from O (no contacts) to 1 (maximal contacts) based on simulations performed with the MHAS21680UT structure (a)-(e) or the MHAS2168IN homology model (f)-(j)- The lipid color code is the same as in Fig. 3a. The transmembrane helices of MHAS2168 are numbered and depicted as rainbow-colored bars. Linker helices A and B are indicated with yellow arrows. The red dotted lineisthe threshold above which thecontact has been considered as relevant.

**Extended Data Figure 8.**
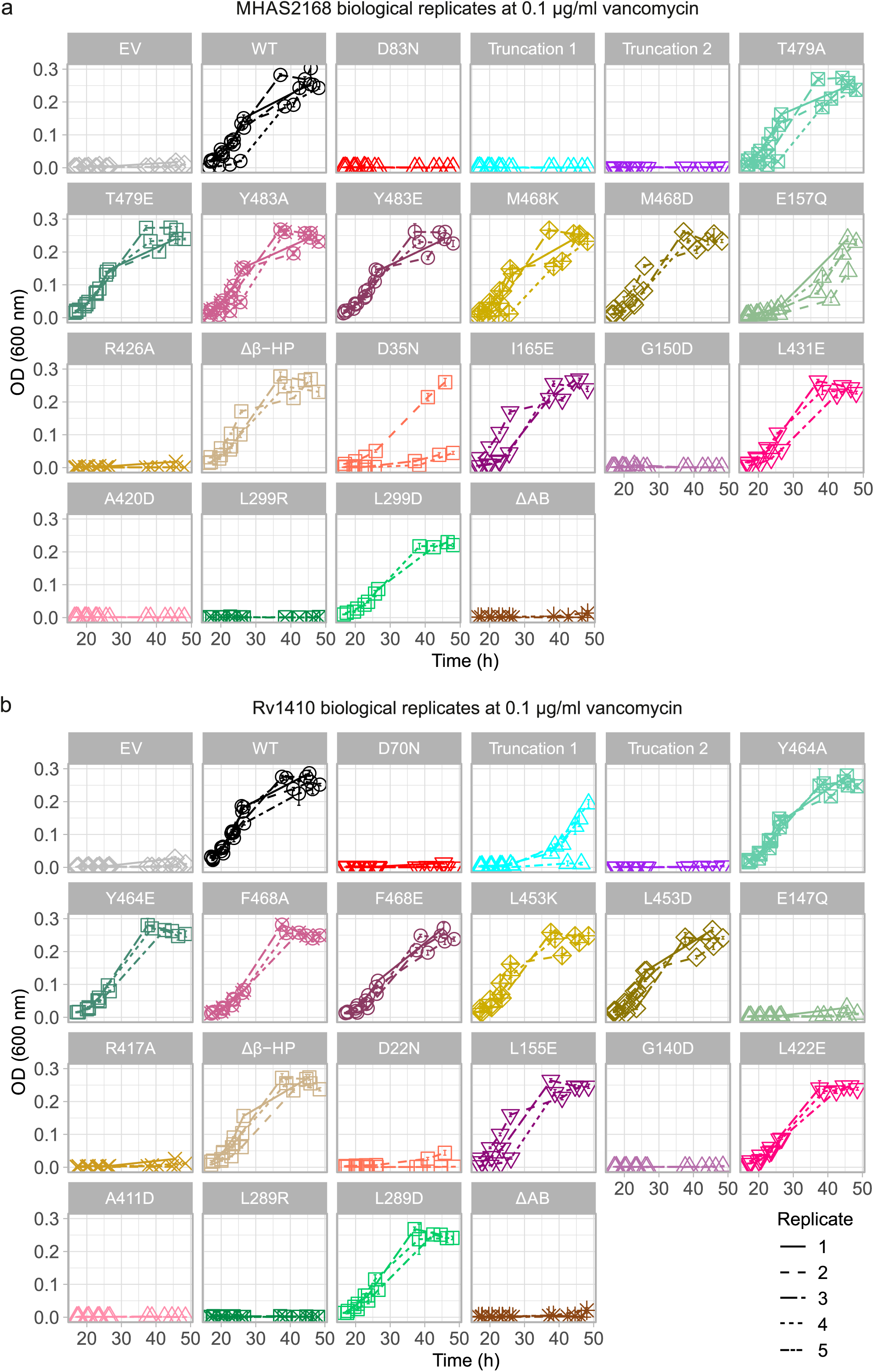
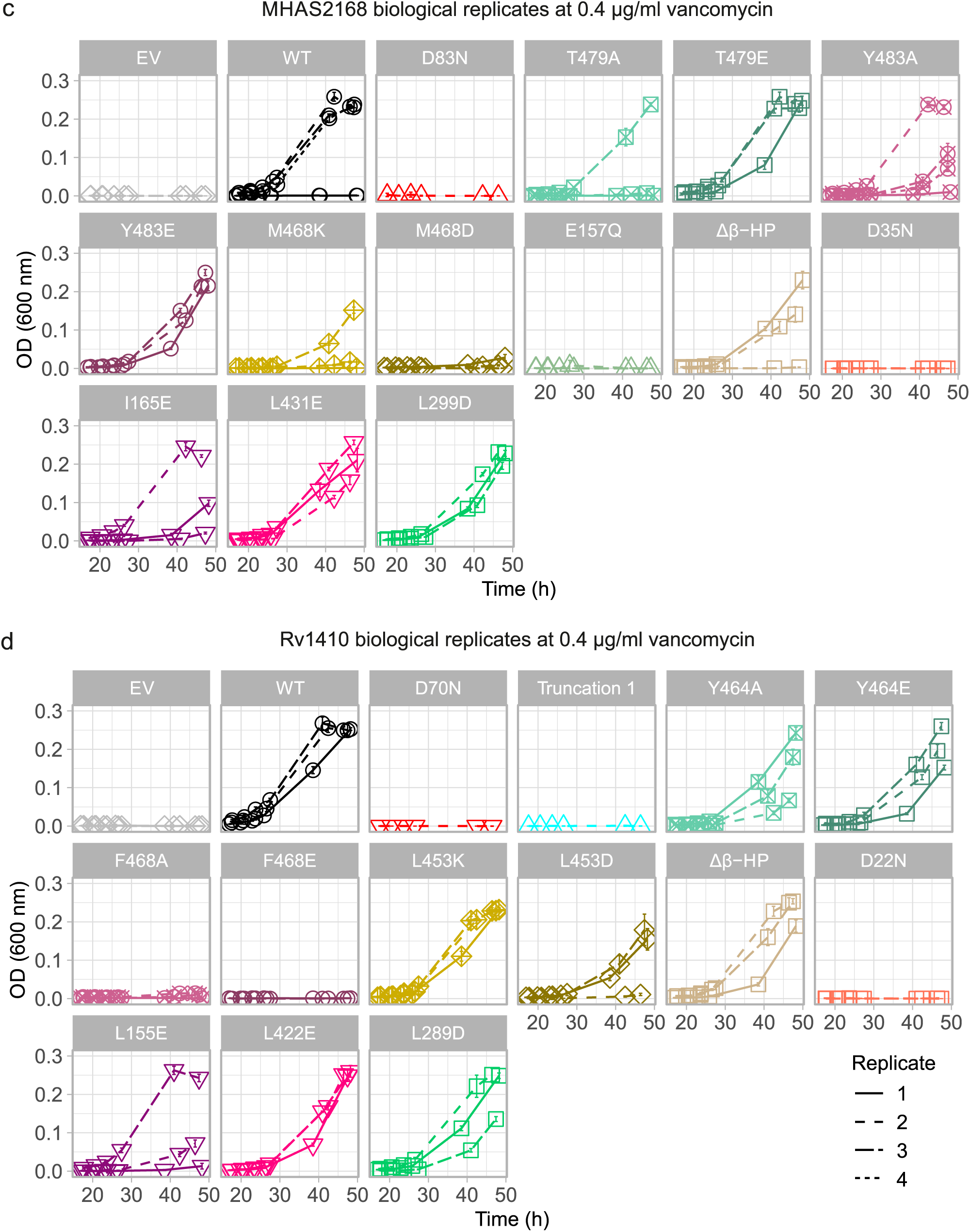
All vancomycin sensitivity assay results at 0.1 and 0.4 µg/ml vancomycin concentration. Vancomycin sensitivity assays in *M. smegmatis* **dK** O cells, complemented with empty vector control (EV), wild type LprG/Rv1410 or MHAS2167/68 operon (WT), or mutant operons where LprG (Rv1411/MHAS2167) is intact, but the transporter Rv1410/MHAS2168 exhibits mutations as indicated. For each tested mutant, growth curves from all biological replicates (3-5) grown at 0.1 µg/ml (a)-(b) or 0.4 µg/ml (c)­ (d) vancomycin concentration are shown. The error bars of the growth curves denote the standard deviation of four technical replicates. (a) and (c) MHAS2168 mutants and controls.

**Extended Data Figure 9.**
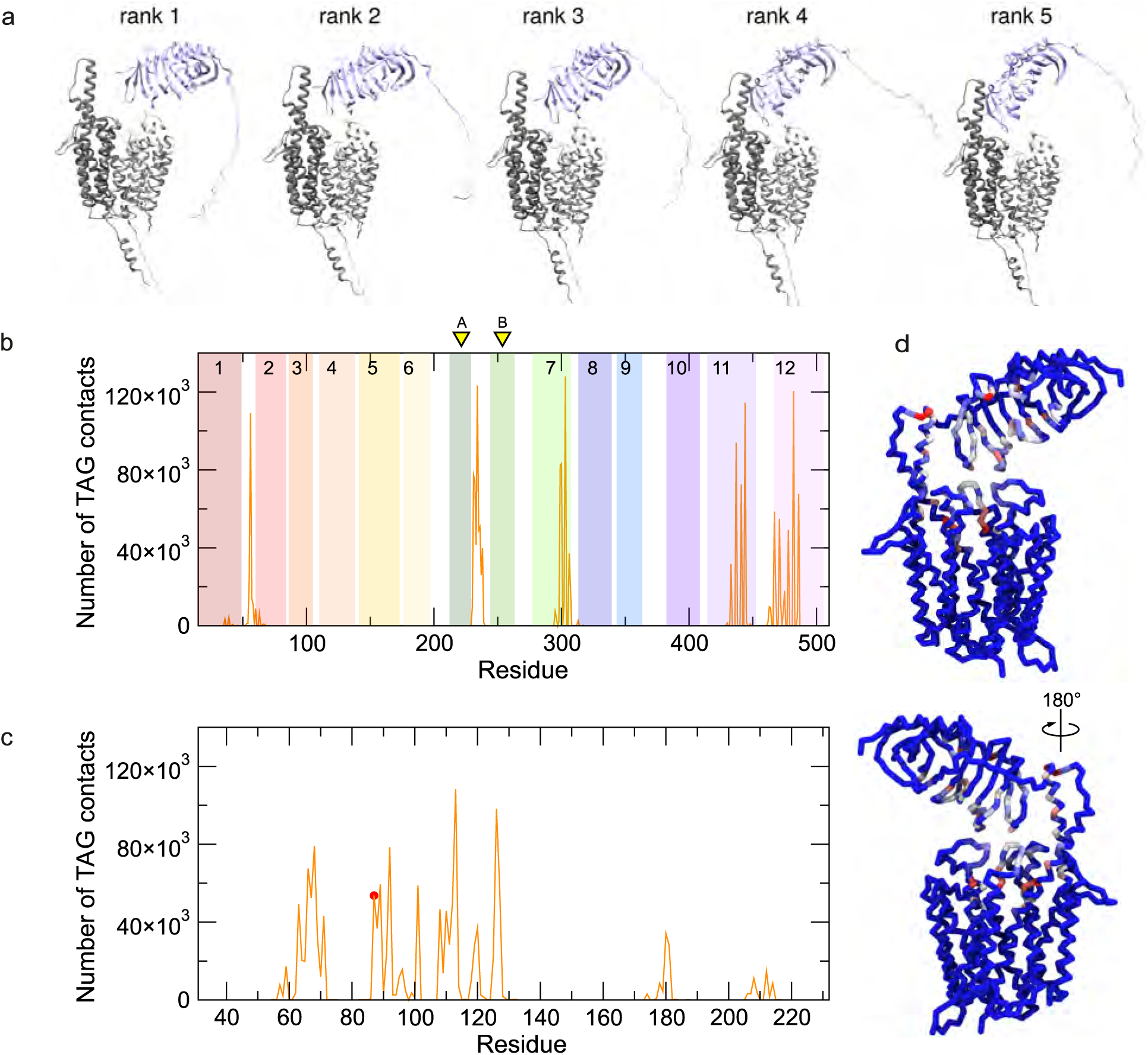
Analysis of TAG contacts with MHAS2168^OUT^-LprG during MD simulations. (a) Five possible models of the MHAS2168^OUT^-LprG complex predicted by Colabfold. The rank 2 model was selected for the MD simulations. (b) Average number of MHAS2168-TAG contacts among the five repeat simulations {orange). The transmembrane helicesof MHAS2168 are numbered. and depicted as rambow-colored bars. Linker helices A and Bareindicated with yellow arrows. (c) LprG-TAG contacts{orange). T87, which corresponds to V91 in M. tuberculosis LprG, is indicated in red.{d) TAG contacts projected onto the protein backboneand coloredfromblue(nocontacts)tored (largenumberof contacts).

**Extended Data Video 1. Video of TAG transfer between MHAS2168 and LprG.** The video shows coarse-grained MD simulation of TAG transfer between MHAS2168 and LprG depicted on Figure 6a, replicate 1. MHAS2168 N-domain, light gray; MHAS2168 C-domain, dark gray; LprG, pale lilac. For clarity, the three TAG acyl tails have been colored differently (yellow, green, and purple).

## Supplementary tables

**Table S1.** X-ray data collection and refinement statistics. In parentheses, parameters of the highest resolution shell are shown.

|  | MHAS2168 + Nb_H2 |  |
| --- | --- | --- |
| Data collection | Crystal I<br>(Full dataset) | Crystals II and III<br>(Merged dataset) |
| Space group | P 1 2<sub>1</sub> 1 | P 1 2<sub>1</sub> 1 |
| Cell dimensions |  |  |
| *a*, *b*, *c* (Å) | 57.75, 160.68, 82.78 | 57.78, 160.7, 82.96 |
| $\alpha$, $\beta$, $\gamma$ (°) | 90.0, 108.854, 90.0 | 90.0, 109.014, 90.0 |
| Resolution (Å) | 2.7 (2.76-2.70) | 2.7 (2.76-2.70) |
| R<sub>meas</sub> (%) | 6.2 (124.1) | 10.7 (183.8) |
| I/$\sigma$ (I) | 11.44 (0.97) | 8.77 (0.97) |
| Completeness (%) | 94.9 (97.8) | 99.3 (99.7) |
| cc(1/2) | 99.9 (46.7) | 99.9 (34.7) |
| Refinement |  |  |
| Resolution range (Å) |  | 44.7 - 2.7 (2.797 - 2.7) |
| No. unique reflections |  | 39031 (3911) |
| R-work/R-free |  | 0.2450/0.2915 |
| No. of atoms |  |  |
| Macromolecules |  | 8824 |
| Ligands |  | 0 |
| Solvent |  | 0 |
| Average B-factor |  | 92.85 |
| RMS deviations |  |  |
| Bonds |  | 0.003 |
| Angles |  | 0.73 |
| Ramachandran favored (%) |  | 98.29 |
| Ramachandran allowed (%) |  | 1.71 |
| Ramachandran outliers (%) |  | 0 |
| Rotamer outliers (%) |  | 0.23 |
| Clashscore |  | 2.00 |
| RSRZ (%) |  | 14.4 |
| MolProbity score |  | 0.77 |

**Table S2.** Cryo-EM data collection and processing statistics.

|  | **MHAS2168 + Mb_H2** | **Rv1410 + Mb_F7** |
| --- | --- | --- |
| **Data collection and processing** |  |  |
| Magnification | 130 000x | 130 000x |
| Voltage (kV) | 300 | 300 |
| Electron exposure (e-/Å<sup>2</sup>) | 66.54 | 65.0 / 55.0 |
| Defocus range (µm) | -1 to -2.5 µm | -1 to -2.5 µm |
| Pixel size (Å) | 0.325 (in super-resolution)<br>0.65 (in construction) | 0.325 (in super-resolution)<br>0.65 (in construction) |
| Symmetry imposed | C1 | C1 |
| No. of initial particle images | 4 759 395 | 1 833 683 |
| No. of final particle images | 402 229 | 127 196 |
| Map resolution (Å) | 3.99 | 7.51 |
| FSC threshold | 0.143 | 0.143 |

**Table S3.** MD simulations. 1-palmitoyl-2-oleyl-phosphatidylethanolamine, POPE; 1-palmitoyl-2-oleyl-phosphatidylglycerol, POPG; cardiolipin, CDL; 1-palmitoyl-2-oleyl-phosphatidylinositol, POPI; triacylglycerol, TAG. ul = upper leaflet; ll = lower leaflet. **^‡^** = The systems with 2 and 1 TAG hydrophobic tails pointing upwards (towards LprG) are identical.

| System | Protein particles | Lipids | Membrane composition (leaflet %) | Water particles | Ions (Na<sup>+</sup>, Cl<sup>-</sup>) | Total particles | Box size (nm<sup>3</sup>) |
| --- | --- | --- | --- | --- | --- | --- | --- |
| MHAS2168<sup>OUT</sup> | 985 | ul: 100 molecules<br>ll: 100 molecules<br>2942 particles | POPE: 34<br>POPG: 29<br>CDL: 15<br>POPI: 21<br>TAG: 1 | 5631 | 222, 63 | 9843 | ca. 1137 |
| MHAS2168<sup>IN</sup> | 985 | ul: 100 molecules<br>ll: 98 molecules<br>2918 particles | POPE: 34<br>POPG: 29<br>CDL: 15<br>POPI: 21<br>TAG: 1 | 5850 | 224, 66 | 10043 | ca. 1166 |
| MHAS2168<sup>IN-TAG</sup> | 985 | ul: 100 molecules<br>ll: 98 molecules<br>2918 particles | POPE: 34<br>POPG: 29<br>CDL: 15<br>POPI: 21<br>TAG: 1 | 5850 | 224, 66 | 10043 | ca. 1166 |
| MHAS2168<sup>OUT-LprG</sup>‡ | 1384 | ul: 196 molecules<br>ll: 189 molecules<br>5649 particles | POPE: 34<br>POPG: 29<br>CDL: 15<br>POPI: 21<br>TAG: 1 | 8798 | 493, 169 | 16493 | ca. 1872 |
| Myco<sup>mem</sup> | - | ul: 141 molecules<br>ll: 141 molecules<br>4142 particles | POPE: 34<br>POPG: 29<br>CDL: 15<br>POPI: 21<br>TAG: 1 | 5702 | 329, 103 | 10276 | ca. 1175 |

**Table S4.**
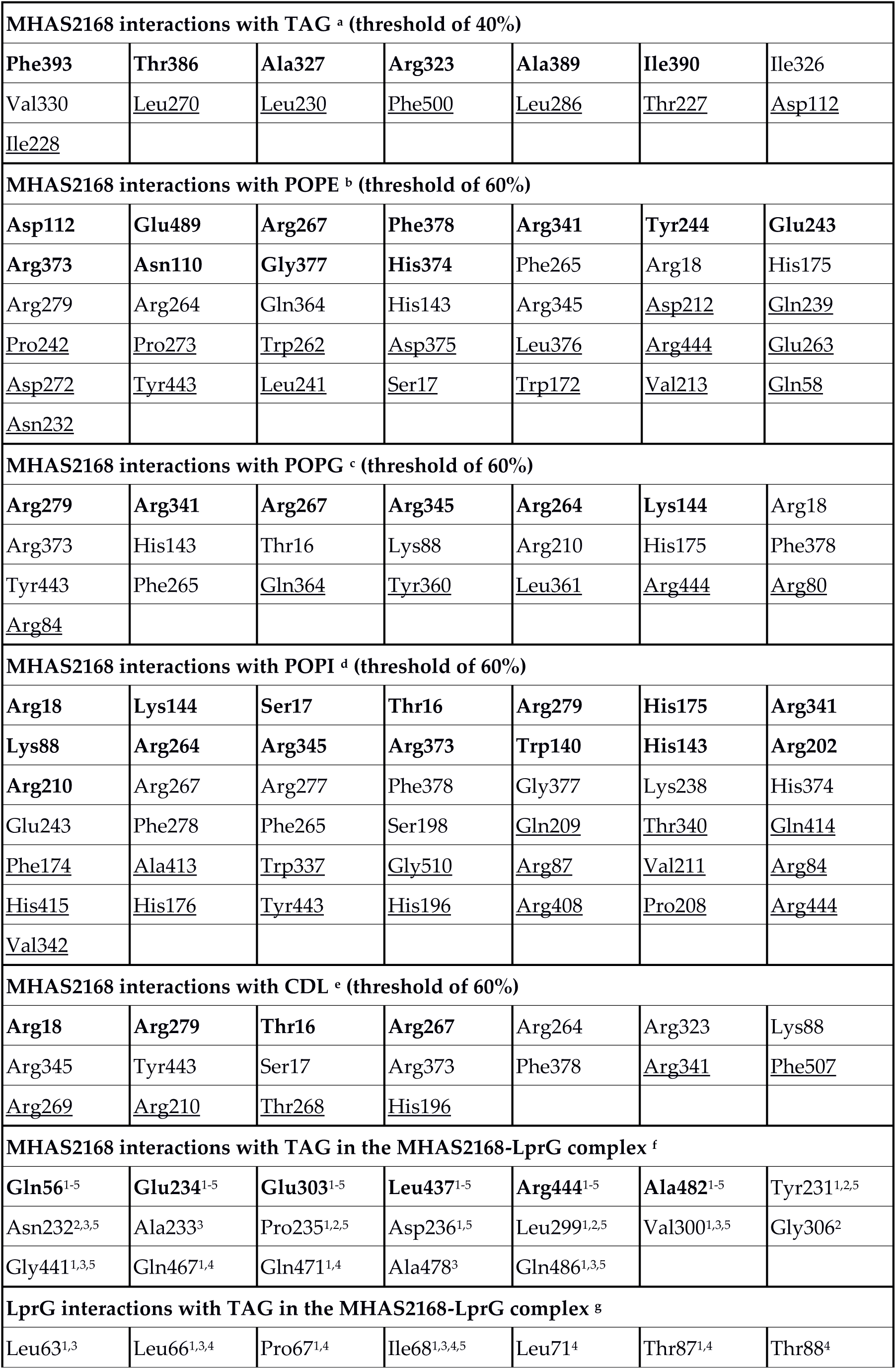
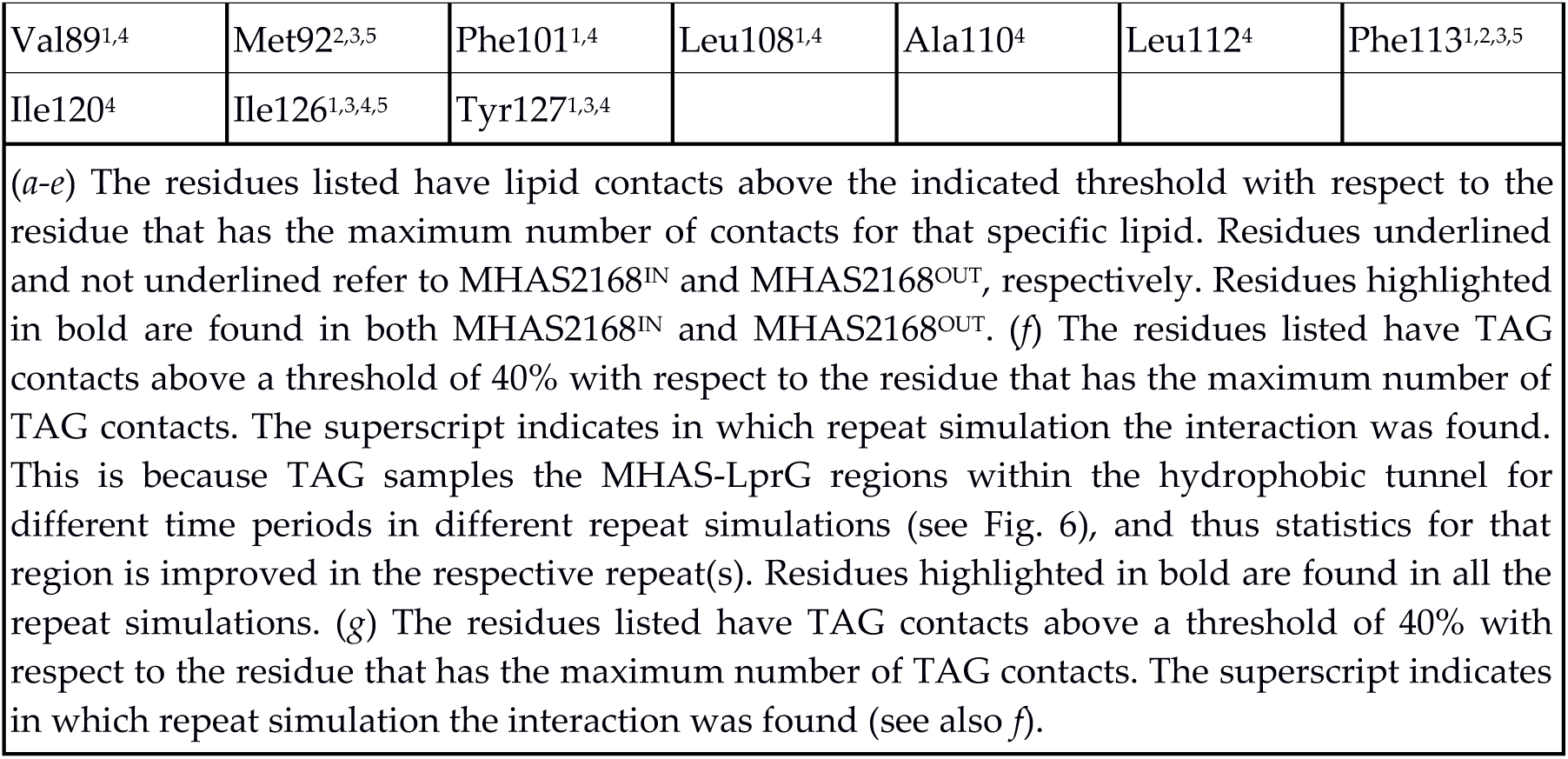
Protein-lipid interactions in CG-MD simulations.

**Table S5.**
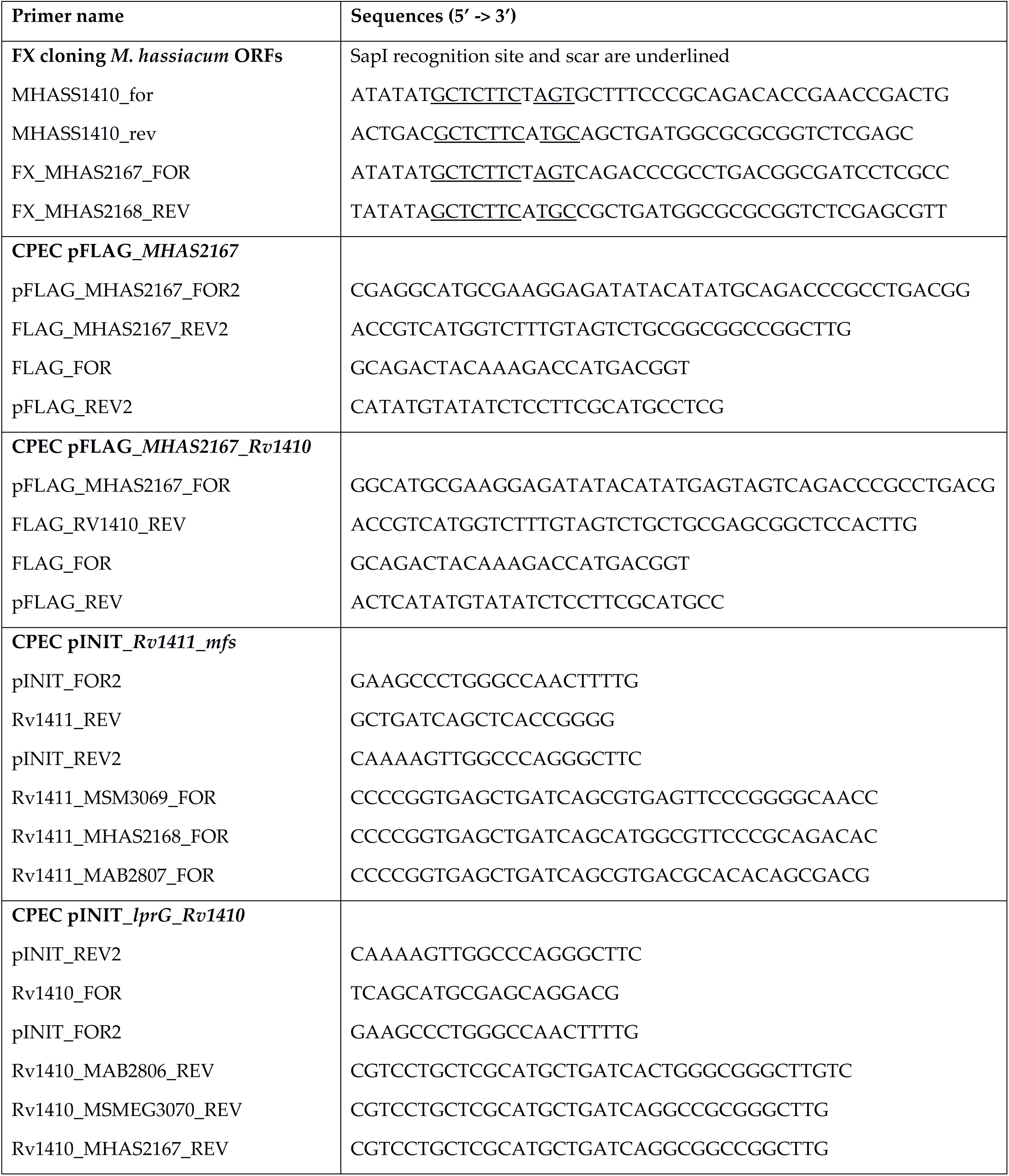
Primers used for generating pFLAG plasmids with shuffled operons.

**Table S6.**
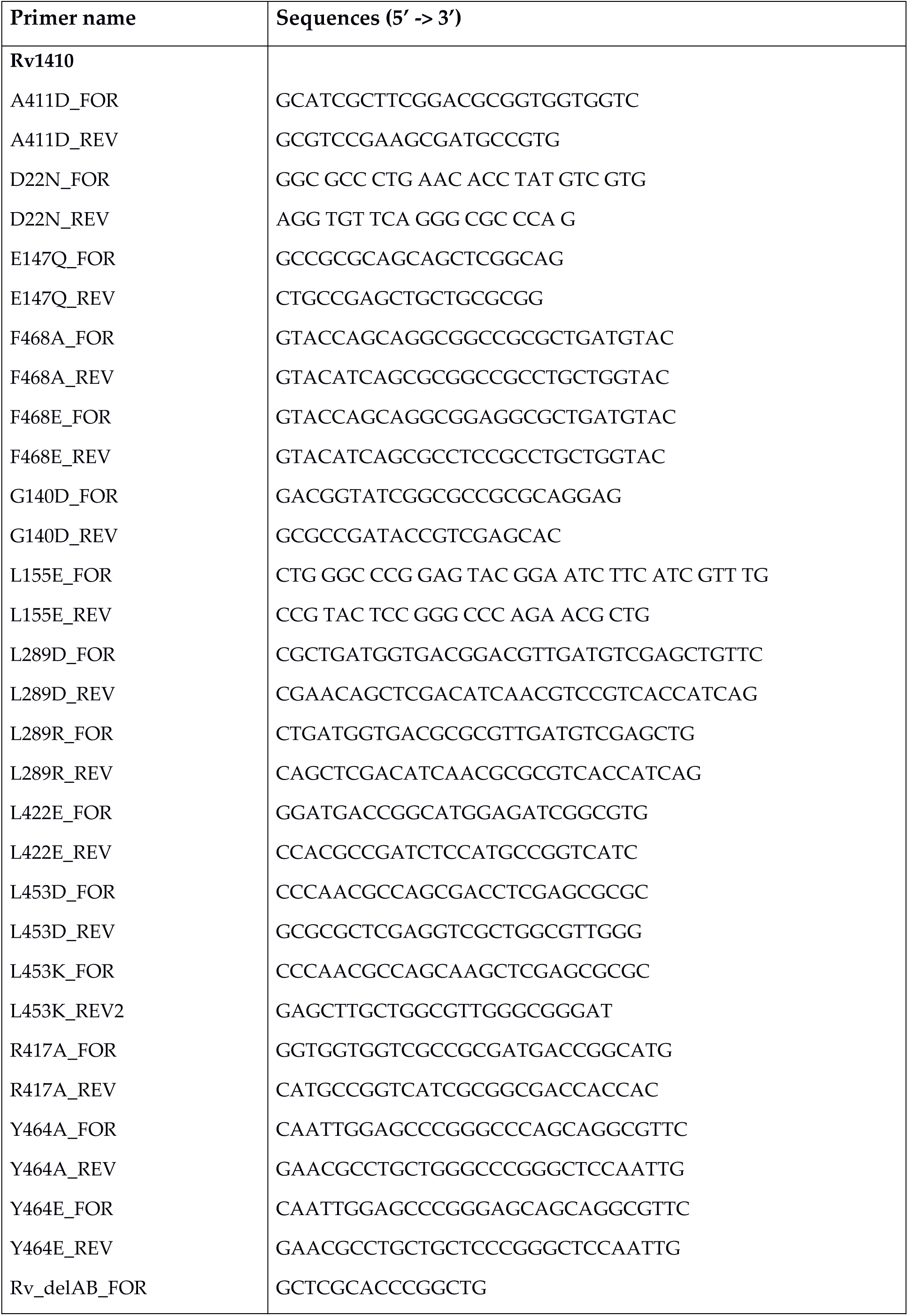
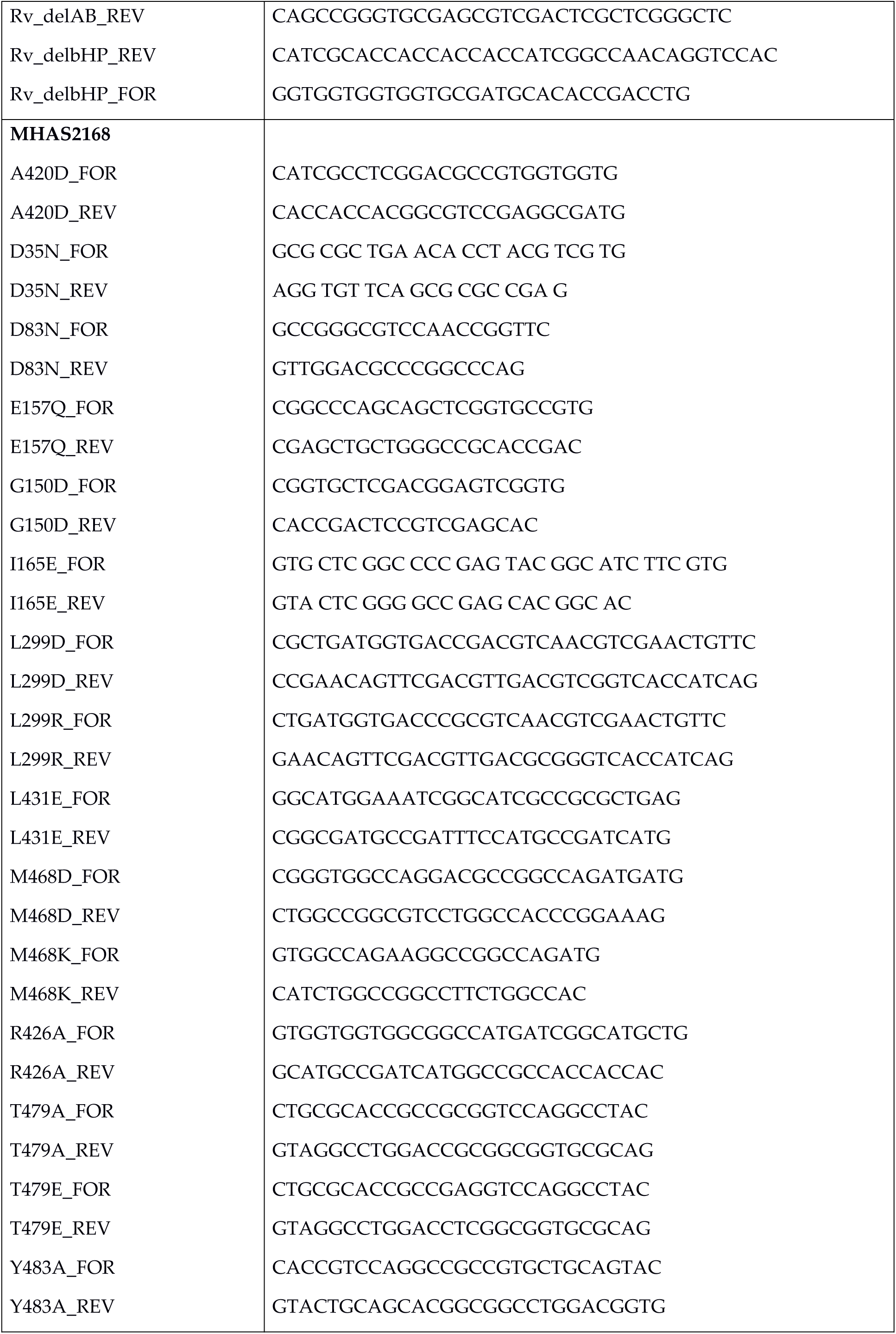
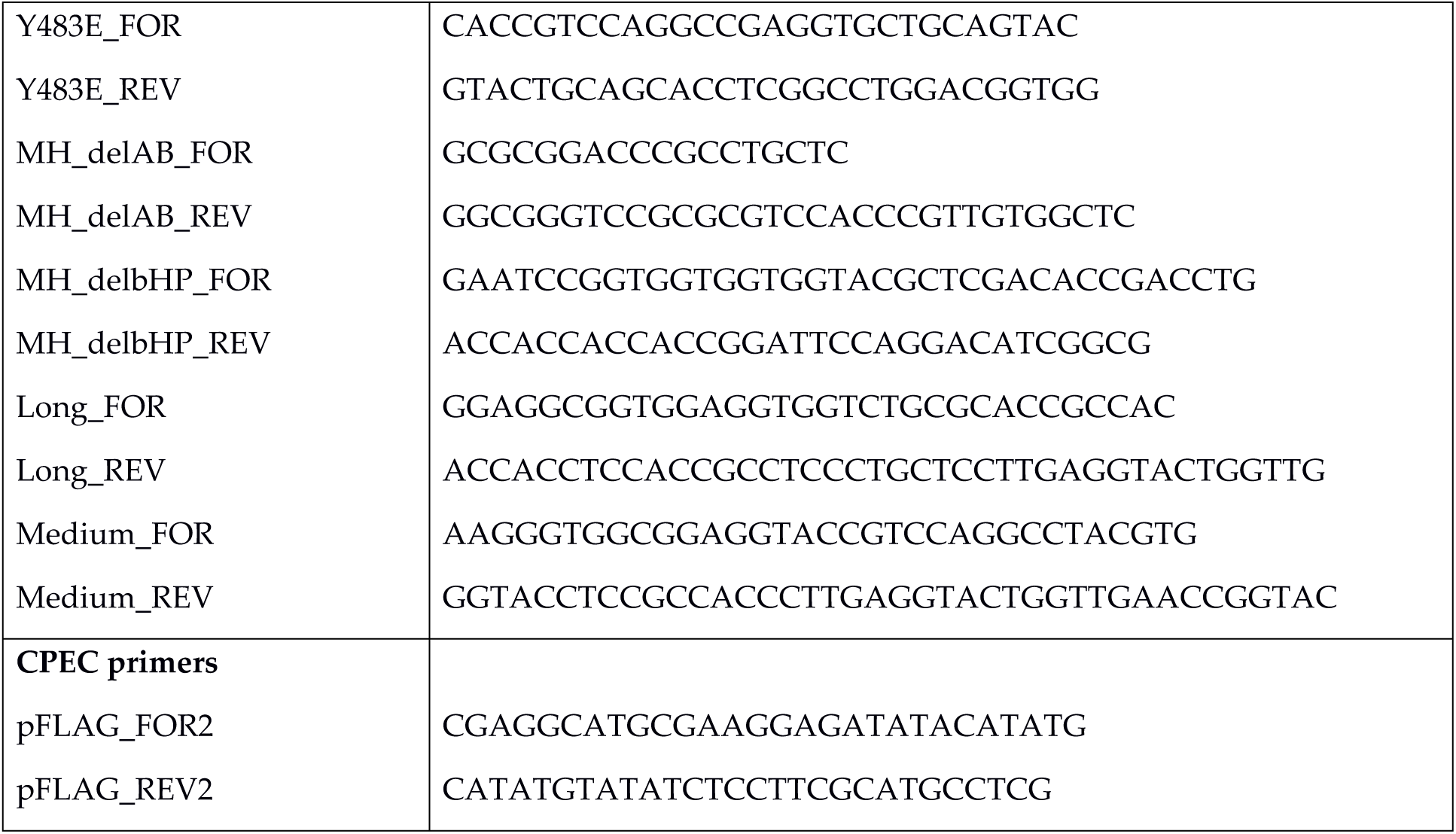
Primers used for introducing mutations into Rv1410 and MHAS2168.

## Supplementary Information

### Rationale for mutant design

The tested mutations were always introduced to both Rv1410 and MHAS2168 to assure the relevance of the phenotype. The mutation sites were chosen according to our MHAS2168^OUT^ crystal structure or the MHAS2168^IN^ homology model, but the mutation locations were checked on Rv1410 structure predictions by ColabFold. Also, conservation in mycobacterial homologues in general (Extended Data Fig. 5) was taken into account. Production of each mutant was tested by Western blotting (Extended Data Fig. 4), to ensure that the phenotypes are not due to insufficient protein production or aberrant folding.

#### Unique ion lock mutations

The only acidic residue that could be protonated/deprotonated during transport cycle in the central cavity is E147_Mtb_/E157_MH_. This glutamate is fully conserved in 17 Rv1410 homologue proteins (Extended Data Fig. 5) and forms a salt bridge with a fully conserved arginine (R417_Mtb_/R426_MH_). This ion lock seems to be unique, as it is not commonly found in other MFS transporters (Extended Data Fig. 6). To test whether the protonation/deprotonation of the glutamate is important for transport, we introduced mutations E147Q_Mtb_/E157Q_MH_ to Rv1410 and MHAS2168, correspondingly. The glutamine cannot be deprotonated. In addition, we introduced the R417A_Mtb_/R426A_MH_ mutations to Rv1410 and MHAS2168, correspondingly, to investigate whether the formation of the ion lock is important for the transporter’s activity.

#### D22/D35

In a paper by Farrow and Rubin^1^, the D22 residue of Rv1410 was investigated as it was speculated to belong to a conserved motif D1. It was discovered that mutation D22A was as sensitive to ethidium as Rv1410 deletion mutant while D22E which also harbours a carboxylate group retained some of ethidium resistance, although not at wild-type level. We reasoned that if the importance of D22 lies in coupling proton translocation to substrate transport, D22N mutation should inactivate Rv1410. Therefore, we introduced the D22N_Mtb_ and D35N_MH_ mutations to Rv1410 and MHAS2168, correspondingly. This aspartate was fully conserved in 17 Rv1410 homologue proteins (Extended Data Fig. 5).

#### β-hairpin truncation

To assess whether the extracellular β-hairpin found between TM9 and TM10 has any impact on the transporter’s function, we decided to truncate the β-hairpin. To do that, we removed the two β-sheets forming the hairpin and residues connecting them, replacing them with a linker formed of four glycine residues. Therefore, Δβ-HP_MH_ is a MHAS2168ΔR373-P382::GGGG mutant and Δβ-HP_Mtb_ is Rv1410ΔR363-P373::GGGG mutant.

#### Truncation of linker helices

To assess whether the linker helices TMA and TMB have any impact on the transporter’s function, we decided to truncate these linker helices to turn Rv1410/MHAS2168 into a classical 12-helix MFS transporter. In this case, the lateral opening between TM2-TM11 in outward-facing conformation is not blocked by linker helices anymore. To achieve that, we deleted the linker helices TMA and TMB and the periplasmic loop connecting them. Since we deemed the remaining cytoplasmic loops to be long enough to connect N- and C-domains, we did not add any extra linker. Therefore, ΔAB_MH_ is a MHAS2168ΔV213-F265 mutant and ΔAB_Mtb_ is Rv1410ΔL203-F255 mutant.

#### Mutations in lateral clefts

To investigate whether lateral openings between N- and C-domain could serve as the entry or exit points for TAGs, we adopted a mutation strategy similar to one used in assessment of MFSD2A^2^. However, we decided to introduce mutations to each lateral cleft, forming in both outward-facing and inward-facing conformations, assessed in our MHAS2168^OUT^ crystal structure and MHAS2168^IN^ homology model. We selected residues on TM2 and TM5 in the middle of each cleft whose side chains (if existing) were faced towards TM11 or TM8, correspondingly. We assumed that if these hydrophobic residues were mutated into glutamates or aspartates, the charged/polar side chains might prevent TAG diffusion through the lateral clefts. To choose whether a glutamate could be introduced into a given location, steric hindrances in each mutant were assessed *in silico* in both MHAS2168^OUT^ and MHAS2168^IN^ conformations. If the side-chain of glutamate seemed to encounter steric hindrances in either of the two conformations, an aspartate was introduced to the loci instead. The mutations are summarized below in Table S7.

**Table S7.** Mutations introduced to lateral openings between N- and C-domains in Rv1410 and MHAS2168.

| Mutation in Rv1410 | Mutation in MHAS2168 | Helices lining the lateral cleft | The lateral cleft is open in |
| --- | --- | --- | --- |
| L155E<sub>Mtb</sub> | I165E<sub>MH</sub> | TM5-TM8 | Outward-facing state |
| G140D<sub>Mtb</sub> | G150D<sub>MH</sub> | TM5-TM8 | Inward-facing state |
| L422E<sub>Mtb</sub> | L431E<sub>MH</sub> | TM2-TM11 | Outward-facing state |
| A411D<sub>Mtb</sub> | A420D<sub>MH</sub> | TM2-TM11 | Inward-facing state |

#### Mutations in cavity

The aim of these mutations was to introduce charge (and bulk) to the hydrophobic wall of the central cavity, in the hope that it might interfere with TAG transport if the molecule resides in the central cavity during its transport. L289_Mtb_/L299_MH_ was chosen as the mutation site because i) its side chain is located in the middle of the hydrophobic central cavity wall in the MHAS2168^OUT^ C-domain, ii) it is fully conserved in 17 Rv1410 homologue proteins (Extended Data Fig. 5), iii) the MHAS2168^IN^ homology model could accommodate a bulky residue in that position. Therefore, L289R_Mtb_/L299R_MH_ mutations were introduced to Rv1410 and MHAS2168, correspondingly, to introduce charge and bulk into the cavity. As a control, L289D_Mtb_/L299D_MH_ mutations were introduced to Rv1410 and MHAS2168, correspondingly, to introduce polar residues with similar size as the original leucine in that position.

#### Truncations of periplasmic helix extensions

In our previous work^3^, we detected a unique “periplasmic loop” between TM11 and TM12 and investigated its truncation mutants which exhibited gradual loss of functionality, the more residues were removed (Long loop = Truncation 1[Δ10aa]; Medium loop = Truncation 2[Δ18aa]; Short loop [Δ26aa]). Our structure showed the presence of TM11 and TM12 helix extensions, instead of a periplasmic loop. When the structures of Rv1410 truncation mutants were predicted by ColabFold platform^4^, surprisingly, the truncation mutants exhibited equal lengths of TM11 and TM12, even if TM12 N-terminal residues had to be conscripted to make up lost length of TM11 C-terminus or vice versa. We decided to design truncation mutants in MHAS2168 to confirm that loss of helix length incurs inactivity of the transporter. However, instead of deleting the corresponding residues of Rv1410 truncation mutants in MHAS2168, we decided to mimic the tertiary structure of these truncations by deleting residues from both TM11, TM12 and the loop connecting them, and introducing a glycine linker (of similar length to the original truncation mutants in Rv1410) between the remaining helices to ensure that the original helices are not disturbed. Therefore, Truncation 1_MH_ is a MHAS2168ΔL453-R474::GGGGGG mutant and Truncation 2_MH_ is MHAS2168ΔE451-A478::GGGG mutant.

#### TM12 mutations

Analysis of periplasmic helix extensions uncovered some conserved features that were common in all the Rv1410 mycobacterial homologues investigated by MSA (Extended Data Fig. 5) and ColabFold structure predictions (Extended Data Fig. 2). The residues on the tip of the TM12 extension (1^st^ α-turn) directed towards the cavity are hydrophobic and aromatic residues are commonly found on TM12 above the cavity (whether on the 4^th^ or 5^th^ α-turn or both). Therefore, we decided to mutate these features to assess whether they might play a role in TAG transport. To judge the necessity for aromatic residues to TAG transport, we introduced alanine mutations Y464A_Mtb_, F468A_Mtb_, T479A_MH_, Y483A_MH_ to the sites of aromatic residues above cavity. To assess whether charged and bulkier side-chains at these locations might interfere with TAG transport, we introduced glutamate mutations Y464E_Mtb_, F468E_Mtb_, T479E_MH_, Y483E_MH_ to the aromatic residues above cavity. To investigate whether the hydrophobic tip of TM12 has a role in TAG transport, we introduced mutations with either positive (L453K_Mtb_, M468K_MH_) or negative charge (L453D_Mtb_, M468D_MH_) to this locus.

### Description of MD simulations

#### Lipid interactions in MHAS2168^OUT^ and MHAS2168^IN^

Coarse-grained molecular dynamics (MD) simulations of MHAS2168 were performed in both outward-open (MHAS2168^OUT^) and inward-open conformations (MHAS2168^IN^) embedded in a phospholipid bilayer doped with TAGs, mimicking the mycobacterial plasma membrane (see methods; Table S3; Fig. 3a). The MFS fold and the sequence similarity shared with other transporters from the same family, allowed the generation of a structural model of MHAS2168 in the inward open conformation, using PepT_So2_ as template (see methods).

The embedment of the TAGs in the hydrophobic core of the membrane (Fig. 3b) enables them to probe a range of different positions along the transporter TM helices (Fig. 3c). TAGs visit both lateral openings of the transporter with a preference for TM8 and TM10 (Extended Data Fig. 7). Protein-lipid contact analyses show that TAGs are more strongly interacting with MHAS2168^IN^ (Extended Data Fig. 7; Table S4), with a particular involvement of the TM4 N-terminus, TM7 N-terminus, TM12 C-terminus and the linker helices A and B.

Concerning the interactions of the transporter with the phospholipids, it is found that MHAS2168 strongly interacts with CDL and POPI lipids, which form an annular lipid belt around the transporter that shields it from other membrane components. Small differences are observed between the MHAS2168^OUT^ and MHAS2168^IN^ conformations (Extended Data Fig. 7; Table S4). Different phospholipid interactions were found especially for the connecting loop between TM10 and TM11 (residues 409-415) and for the TM11 C-term (Extended Data Fig. 7), where MHAS2168^IN^ was found to interact more with POPI than MHAS2168^OUT^.

#### TAGs in MHAS2168^OUT^ and MHAS2168^IN^ central cavities

We further investigated the central cavity and its interactions with TAG during the MD simulations of both conformations. The outward-open conformation cavity volume was estimated to be ∼2500 Å^3^, while the inward-open homology model displays a slightly larger cavity and a broader distribution, centred around 2800 Å^3^ (Fig. S1).

**Figure S1.**
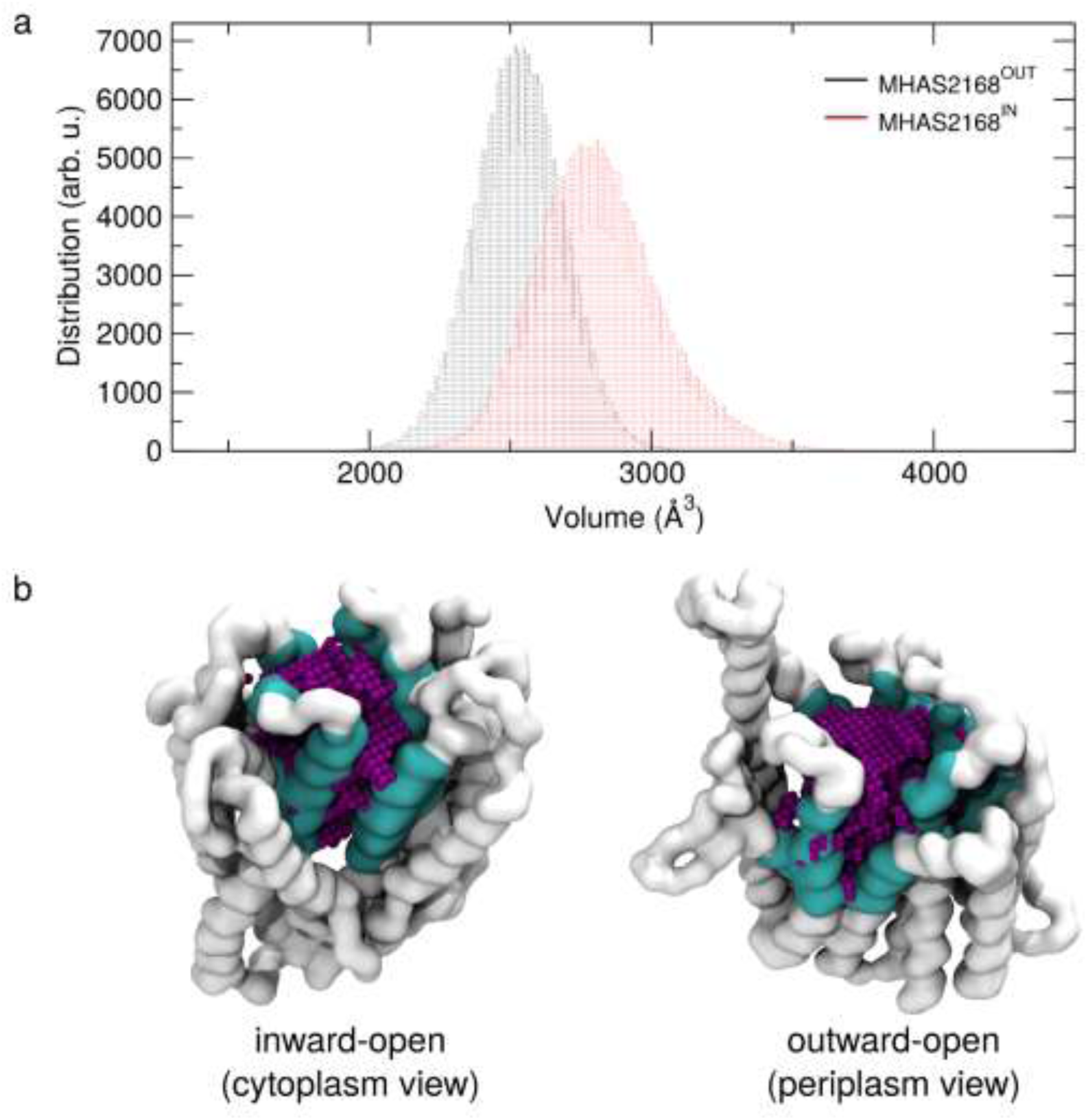
The main cavity of MHAS2168. (**a**) The histograms show the volume of the main cavity encompassed by the N- and C-terminal domains calculated during the MD simulations for the MHAS2168^OUT^ and the MHAS2168^IN^ conformations. (**b**) Snapshots from the MD simulations with the main cavity and the protein backbone atoms highlighted in purple and white, respectively. Residues in cyan were selected for the cavity calculations (see methods).

Furthermore, we recorded the entrance events of the different lipid species into the main cavity of MHAS2168 during MD simulations (Fig. S2). Apart from a few snorkelling events, in which a TAG molecule probes the periphery of the cavity but does not fully enter, the MHAS2168^OUT^ cavity is little accessed by phospholipids and TAGs. Conversely, for MHAS2168^IN^ multiple entry and exit events were observed for POPG, POPE and especially POPI (Fig. S2), in line with our above results according to which POPI creates an annular belt around the protein. Phospholipid entrance and exit events occur via both lateral openings (i.e., TM5-TM8 and TM2-TM11). However, no TAG molecule entered the cavity in the MD simulations, likely because of the much lower TAG concentration compared to the competing phospholipid species. To probe whether the central cavity can accommodate TAG, we ran five additional MD simulations for MHAS2168^IN^, where we inserted one TAG molecule into the main cavity from the beginning of the simulations (MHAS2168^IN-TAG^, see Table S3, Fig. S3). The TAG residence times within the cavity in these 5 simulations are 20, 15, 18, 8 and 74 μs, respectively, enabling us to collect a total of 135 μs of simulation time that allow one to map the TAG interactions within the cavity (Fig. S3). Especially the TM5 N-terminus and the TM8 C-terminus, where the non-proteinaceous density was found in the cryo-EM structure (Fig. 1a), preferably interact with TAG in the simulations (Fig. S3).

**Figure S2.**
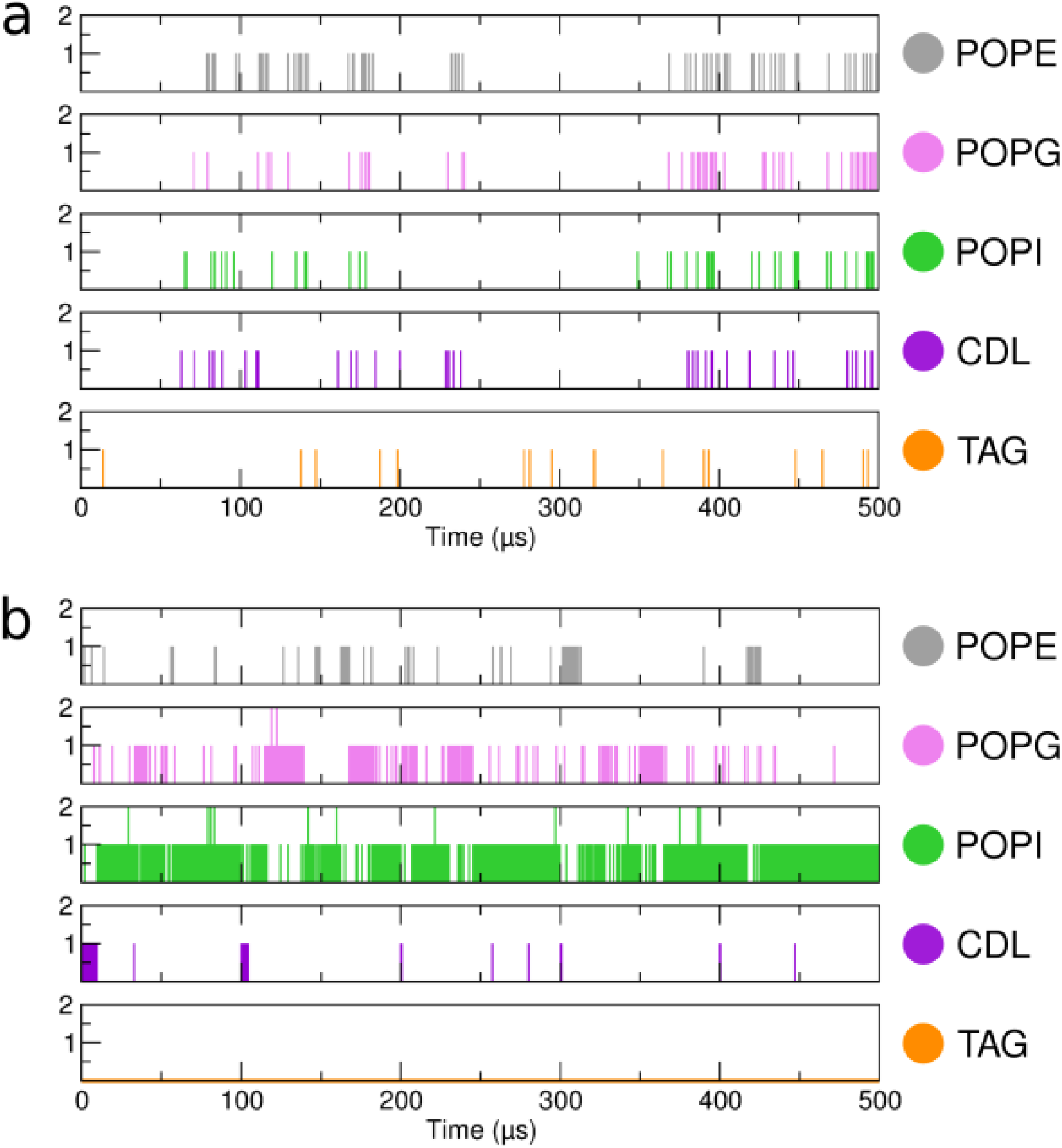
Mapping TAG and phospholipid entrance from the membrane into the transporter during the MD simulations of MHAS2168. Number of entrance and snorkeling events of the membrane components within the protein main cavity during the MD simulation of the MHAS2168^OUT^ (**a**) and MHAS2168^IN^ (**b**) systems. An event of entrance/snorkeling was recorded if the phosphate bead or the TAG backbone bead overlap with the cavity volume shown in Fig. S1. All the five independent repeats of 100 μs each were concatenated together for clarity.

**Figure S3.**
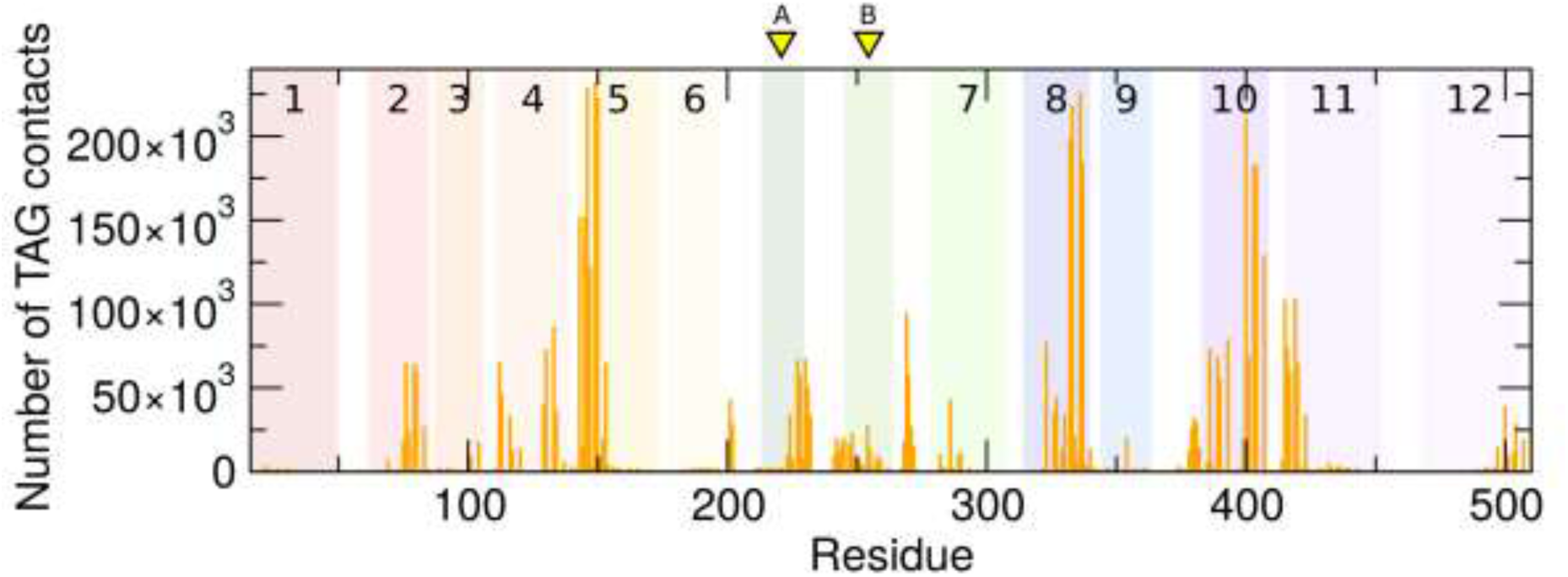
MD simulations of inward-open MHAS2168 with TAG initially placed within the main cavity. Overall protein-TAG contacts (orange) in the 135 μs of collected simulation time of MHAS2168^IN^ (see text). The transmembrane helices of MHAS2168 are numbered and depicted as rainbow-colored bars. Linker helices A and B are indicated with yellow arrows.

In addition, we studied TAG interactions with the central cavity of MHAS2168^OUT^ and its loading process from MHAS2168 to LprG (MHAS2168^OUT^-LprG simulation system, Table S3). To achieve that, a complex of MHAS2168^OUT^ and LprG was built, utilizing the ColabFold^4^ platform. In these simulations, TAG moved between the hydrophobic cavities of MHAS2168^OUT^ and LprG (see description in main text; Fig. 6; Extended Data Fig. 9; Extended Data Video 1).

Initially, the docked TAG was inserted into the transporter with 2 hydrophobic tails pointing upwards (towards LprG) and 1 tail downwards (embedded into the main cavity of MHAS2168, Fig. 6). To probe whether this particular initial configuration could bias the movement of TAG towards LprG, an additional control simulation was carried out in which the TAG molecule was docked with only 1 tail pointing upwards and 2 tails downwards (Fig. S4). In this control simulation, the TAG initially remained in its 2-tails-down configuration for around 80 µs, but then spontaneously transitioned to a 2-tails-up configuration until the end of the simulation run at 100 µs (Fig. S4). When this control simulation was extended (Fig. S4, red line) or used as a starting point for additional 5 repeat simulations with different starting velocities (Fig. S4, lines from 1 to 5), the TAG was found to be loaded into LprG, akin to the independent simulations having a 2-tails-up configuration at the onset of the simulation (Fig. 6). We thus conclude that the initial configuration of the TAG does not determine the outcome of the simulations, and that at least 2 tails of the TAG need to be first oriented towards LprG for the TAG molecule to be spontaneously loaded into LprG.

**Figure S4.**
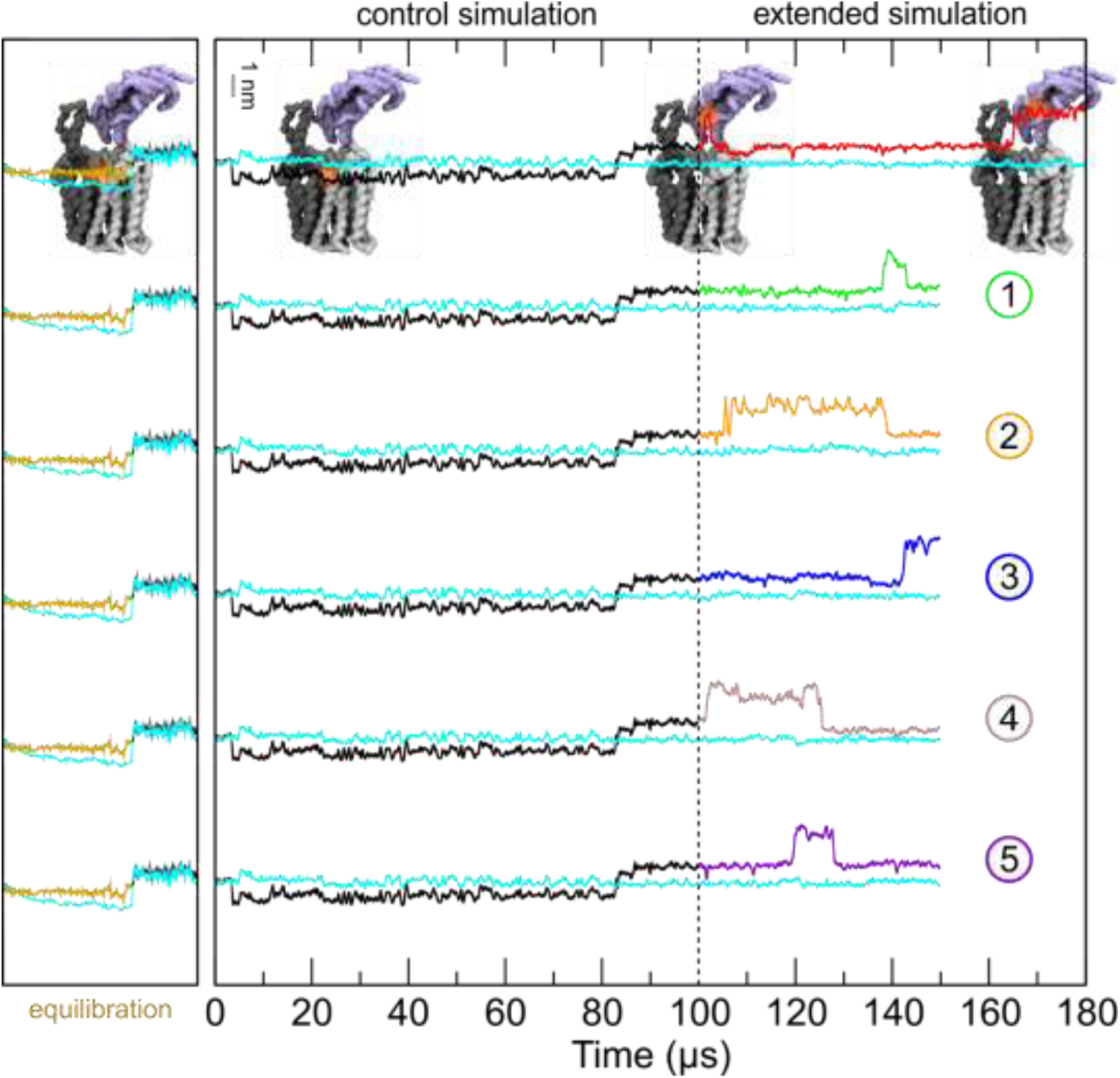
TAG loading into LprG in the control MD simulations of the MHAS2168^OUT^-LprG complex. A control simulation was carried out for 100 µs, starting with TAG in a 2-tails-down configuration. The black line depicts the z-coordinate of the center of mass of the TAG molecule. The cyan line is the z-coordinate of the average center of mass of the phosphate groups of the upper membrane leaflet. The extended simulation (in red), and the five repeat simulations with different starting velocities (green, orange, blue, grey and purple) are indicated and numbered respectively. The gold line is the equilibration phase (see methods). The models of the MHAS2168^OUT^-LprG complex shown in the top panel indicate the position of the TAG (orange shade).

